# Retroelement-Age Clocks: Epigenetic Age Captured by Human Endogenous Retrovirus and LINE-1 DNA methylation states

**DOI:** 10.1101/2023.12.06.570422

**Authors:** Lishomwa C. Ndhlovu, Matthew L. Bendall, Varun Dwaraka, Alina PS Pang, Nicholas Dopkins, Natalia Carreras, Ryan Smith, Douglas F. Nixon, Michael J. Corley

**Affiliations:** Department of Medicine, Division of Infectious Diseases, Weill Cornell Medicine, New York City, New York, USA; TruDiagnostic, Lexington, Kentucky, USA

**Author notes:** Corresponding Author: Michael J. Corley PhD, Assistant Professor, Weill Cornell Medicine, Department of Medicine, Division of Infectious Diseases, New York, NY, USA.

## Abstract

Human endogenous retroviruses (HERVs), the remnants of ancient viral infections embedded within the human genome, and long interspersed nuclear elements 1 (LINE-1), a class of autonomous retrotransposons, are silenced by host epigenetic mechanisms including DNA methylation. The resurrection of particular retroelements has been linked to biological aging. Whether the DNA methylation states of locus specific HERVs and LINEs can be used as a biomarker of chronological age in humans remains unclear. We show that highly predictive epigenetic clocks of chronological age can be constructed from retroelement DNA methylation states in the immune system, across human tissues, and pan-mammalian species. We found retroelement epigenetic clocks were reversed during transient epigenetic reprogramming, accelerated in people living with HIV-1, responsive to antiretroviral therapy, and accurate in estimating long-term culture ages of human brain organoids. Our findings support the hypothesis of epigenetic dysregulation of retroelements as a potential contributor to the biological hallmarks of aging.

## Introduction

Retroelements such as human endogenous retroviruses (HERVs) and long interspersed nuclear elements (LINEs) constitute a significant portion of the human genome(*1*, *2*). While the majority of retroelements embedded within the human genome are typically repressed by epigenetic mechanisms that include DNA methylation and chromatin modifications, the activity of specific HERVs and LINEs in the human genome has been shown to impact gene regulation, gene expression, genomic stability, development, and the pathogenesis of various human diseases(*3–5*). Moreover, the resurrection of particular HERVs and LINEs have been linked to the aging process(*6*– *8*). These studies support a key role of retroelement activity in the biological hallmarks of aging. However, the interplay between the locus-specificity of certain retroelements and aging remains largely unexplored.

Epigenetic clocks are highly accurate biological markers of aging based on patterns of DNA methylation at specific regions of the human genome. They offer a way to measure biological age, which can be distinct from chronological age(*9–13*). However, current first, second, and third-generation epigenetic clocks have been constructed from underlying DNA methylation features that have not focused on retroelements(*9*, *10*, *14–16*). Yet, loss of DNA methylation has been shown to occur with age at repetitive DNA sequences, introns, and intergenic regions of the genome(*17*). Moreover, DNA methylation plays a crucial role in the regulation of retroelements and the reactivation of certain HERVs and LINEs have been observed to increase with age(*6–8*). The potential of using the DNA methylation states of specific HERVs and LINEs in epigenetic clocks for estimating biological age remains unclear.

Here, we investigate DNA methylation dynamics of retroelements as biological age predictors. We enhanced the annotation of Illumina’s MethylationEPIC v1.0 platform to identify CpGs located within manually curated locus-specific HERVs and LINEs(*18*), which were previously unaccounted for in standard and custom add on annotations. Using these retroelement CpGs, three novel epigenetic clocks were developed from DNA methylation data from 12,670 individuals across the lifespan: HERV-Age, based on DNA methylation states of HERVs; LINE-1-Age, based on DNA methylation states of LINEs; and a composite Retroelement-Age clock based on DNA methylation states of HERVs and LINEs. We further constructed a composite Retroelement-Age V2 clock compatible with the MethylationEPIC v2.0 platform and demonstrate that retroelement clocks extend to diverse human tissues and across mammalian species using DNA methylation from GTEx(*19*) and from the Mammalian Methylation Consortium(*20*). Our results substantiate the hypothesis that the dysregulation of retroelements may play a role in the distinctive biological features associated with aging.

## Results

### Annotation of CpGs located in locus-specific HERVs and LINEs

Analysis of MethylationEPIC data relies on a reference annotation of CpGs typically utilizing Illumina’s default annotation that does not identify whether a CpG is located within a locus-specific HERV or LINE(*21*). Moreover, manually curated locus-specific HERV CpGs are not available in enhanced annotations of the MethylationEPIC(*22*, *23*). Hence, we leveraged a manually curated locus-specific HERV annotation of 60 HERV families(*18*) to annotate probes included on the Illumina Infinium MethylationEPIC (EPIC) V1 array. This annotation approach identified that 5,436 probes (0.62% of EPIC) assessed DNA methylation at a CpG within a HERV contained in the Telescope database (**Fig. 1A**). Next, we sought to extend our annotation approach to active LINE elements of the EPIC array. We used Telescope’s locus-specific LINE annotation of 13,545 loci derived from L1Base(*18*) to annotate the EPIC array. This annotation approach identified that 5,543 probes (0.64% of EPIC) assessed DNA methylation at a CpG within a LINE element (**Fig. 1A**). As expected, the EPIC HERV CpGs and LINE CpGs were significantly enriched in regions more than 4kb away from a CpG island referred to as OpenSea and intergenic genomic locations (**Fig. 1B-C**). We observed 1,088 locus-specific HERVs contained more than one CpG measured by the EPIC array (**Fig. 1D**). The greatest coverage on the EPIC array was for HERV HUERSP3_6p21.32 that contained 39 CpGs covered by the MethylationEPIC array (**Fig. 1D**). The greatest coverage of LINE elements on the EPIC array was for L1FLnl_1p31.3z (**Fig. 1E**). We examined the enrichment of Telescope annotated CpGs on the MethylationEPIC using the KnowYourCG tool in SeSAme(*23*) and databases for consensus transcription factor binding sites (TFBSs), ENCODE chromHMM states(*24*), consensus histone modifications(*24*), CpG island shores (up to 2kb between CpG islands), CpG island shelves (regions 2 to 4kb between CpG islands) and open seas (rest of the genome), RepeatMasker annotations(*25*), A/B euchromatic and heterochromatic compartments, and CpGs identified in epigenome-wide association studies. We observed that HERV and LINE CpGs were enriched in ATF7IP bindings sites, quiescent and heterochromatin regions, and B2/B3 high DNA density sub-compartments of heterochromatin consisting of nucleolar associating domains and lamina associating domains (**Fig. 1F-H**). ATF7IP has been shown to be a key interaction factor required for silencing retroelements(*26*). Additionally, HERV and LINE CpGs were enriched by the repressive histone modifications H3K9me2 and H3K9me3 known to transcriptionally silence transposable elements(*27*) (**Fig 1I**). As predicted, HERV and LINE CpGs were enriched in LTR, LINE, ERVK, RV1, ERVL, L1 based on RepeatMasker (**Fig. 1J-K**). We also observed a significant enrichment of HERV and LINE CpGs in loci previously associated with sex, smoking, amyloid plaques, weight loss, cancer, and HIV infection (**Fig. 1L**). No significant enrichment was observed in specific CpG island regions (**Fig. 1M**).

**Fig. 1.**
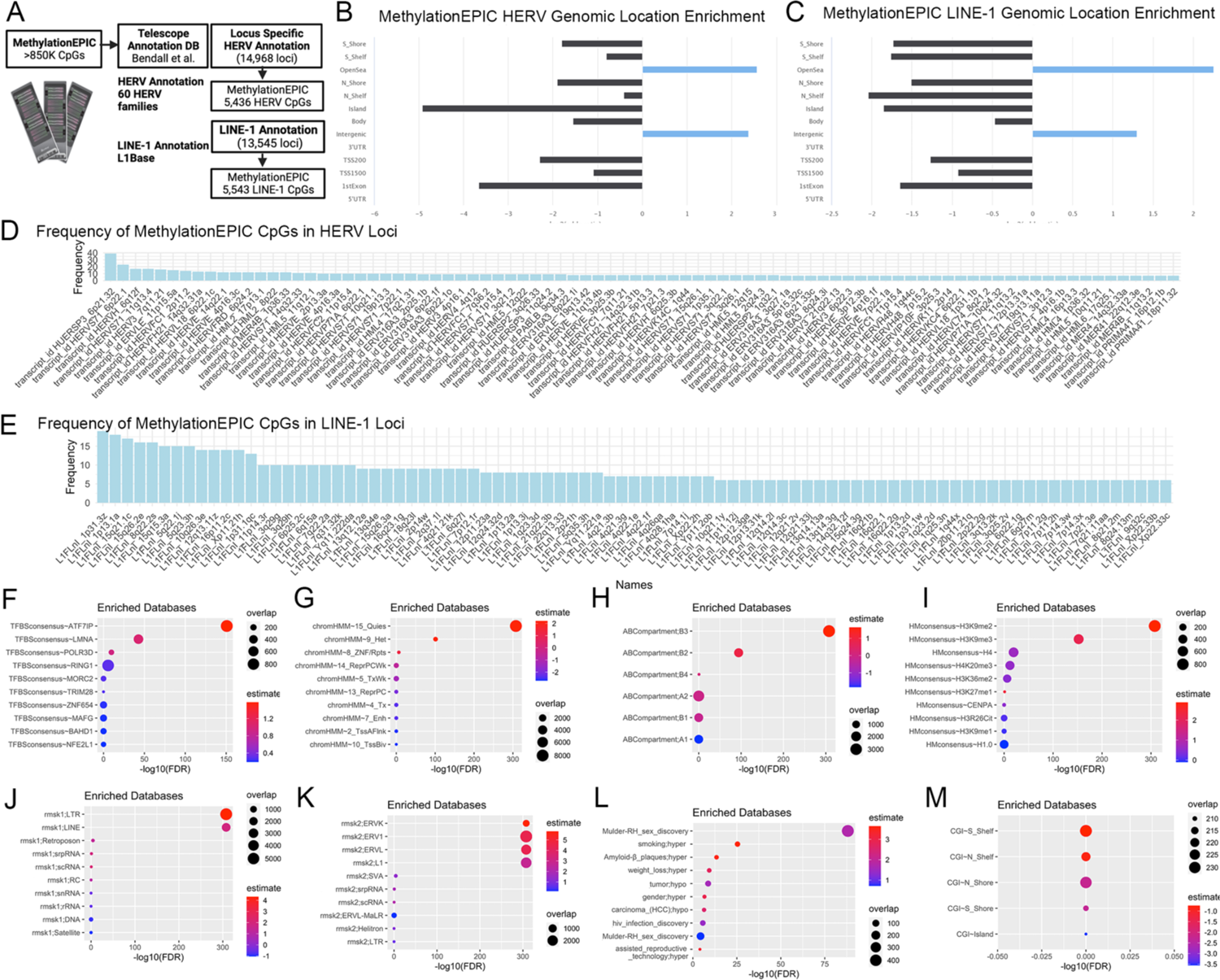
Retroelement Annotation of MethylationEPIC based on Locus Specific HERV and LINE Annotations. **A.** Diagram of Telescope Annotation Database consisting of 14,698 HERV loci and 13,545 LINE-1 elements utilized to annotate HERV and LINE-1 CpGs assayed by the MethylationEPIC v1.0 platform. **B.** Genomic enrichment analysis of 5,436 HERV loci MethylationEPIC CpGs and **C**. 5,543 LINE-1 MethylationEPIC CpGs. **D.** Frequency of CpGs located in Telescope annotated HERV and **E.** LINE elements. **F.** Enrichment plot of Telescope annotated HERV and LINE CpGs in consensus transcription factor binding sites (TFBS) **G.** ENCODE ChromHMM chromatin states, **H.** A/B euchromatic and heterochromatic compartments, **I.** histone modifications, **J-K.** RepeatMasker annotations, **L.** epigenome-wide association CpGs, and **M.** CpG island regions. Fisher’s exact test, estimate represents fold enrichment, and overlap the number of CpGs.

### Development of epigenetic clocks based on Human Endogenous Retroviruses (HERV-Age) and LINE transposable elements (LINE-1-Age) DNA methylation states

Given the role of DNA methylation in regulating HERVs, we first sought to develop an epigenetic clock based on HERV DNA methylation states. We leveraged a dataset of blood DNAm EPIC array data from 12,670 people (40.89% female) with chronological ages ranging from 12 to 100 years old. The database was quality controlled, normalized, and filtered to 5,436 CpGs based on our HERV annotation. The dataset was preprocessed by randomly splitting into an 80% training and 20% test data set. A generalized linear elastic net model was fit with 10-fold cross-validation using glmnet on the 80% training dataset of 10,138 samples (**Fig. 2A**). This approach yielded a new HERV-based epigenetic clock (HERV-Age) based on 954 CpG sites located within a HERV element that performed well in both training (**Fig. 2B**) and testing (**Fig. 2C**) datasets correlating significantly (r=0.93 training and r=0.89 testing) with chronological age (**Table S1**). The median “error” (MAE) defined by the median absolute difference between HERV-Age and chronological age was less than 4 years with 3.12 years in training and 3.8 years in testing datasets. Notably, we observed that 7 CpGs (cg16951108, cg19786602, cg11630939, cg08960549, cg08883146, cg24004238, and cg23987336) that were part of the HERV-Age were located at the HML-2_17p13.1 locus. These CpGs related to a 517bp long intergenic non-coding RNA (lincRNA) ENST0000577807 and overlapped with annotated enhancer regions based on ENCODE ChromHMM. The HERV-K (HML-2) family are the most recently integrated HERV family and recently the activation of HERV-K has been linked to aging and cellular senescence(*6*). Additionally, loci including HERVL18_6p22.1c (cg11341932, cg12212060, cg19203575, cg20850981, cg21177257, cg23702226), HUERSP3_6p21.32 (cg03978169, cg13904130, cg15737123, cg18927185, cg22136013, cg23917817), and MER41_22q12.3e (cg05209515, cg06637550, cg09686907, cg20866592, cg26440042, cg26538224) contained 6 CpGs each that were part of HERV-Age. CpGs in HERVL18_6p22.1c were within the *ZSCAN31* and *ZKSCAN3* genes. *ZKCAN3* is a master transcriptional repressor of autophagy(*28*) and has been shown to mitigate cellular senescence by stabilizing heterochromatin(*29*). CpGs in HUERSP3_6p21.32 were in an intergenic region of heterochromatin/ZNF genes and repeats. CpGs in MER41_22q12.3e were near an enhancer region of the anti-inflammatory and metabolic regulator *C1QTNF6* gene(*30–34*). The MER41 family is an endogenized gammaretrovirus whose LTR has been coopted as an IFNG-inducible binding site bound by STAT1 and/or IRF1 promoting inflammasome production(*35*). These genomic regions implicated in HERV-Age highlight the potential role of HERVs in immune age-related processes.

**Fig. 2.**
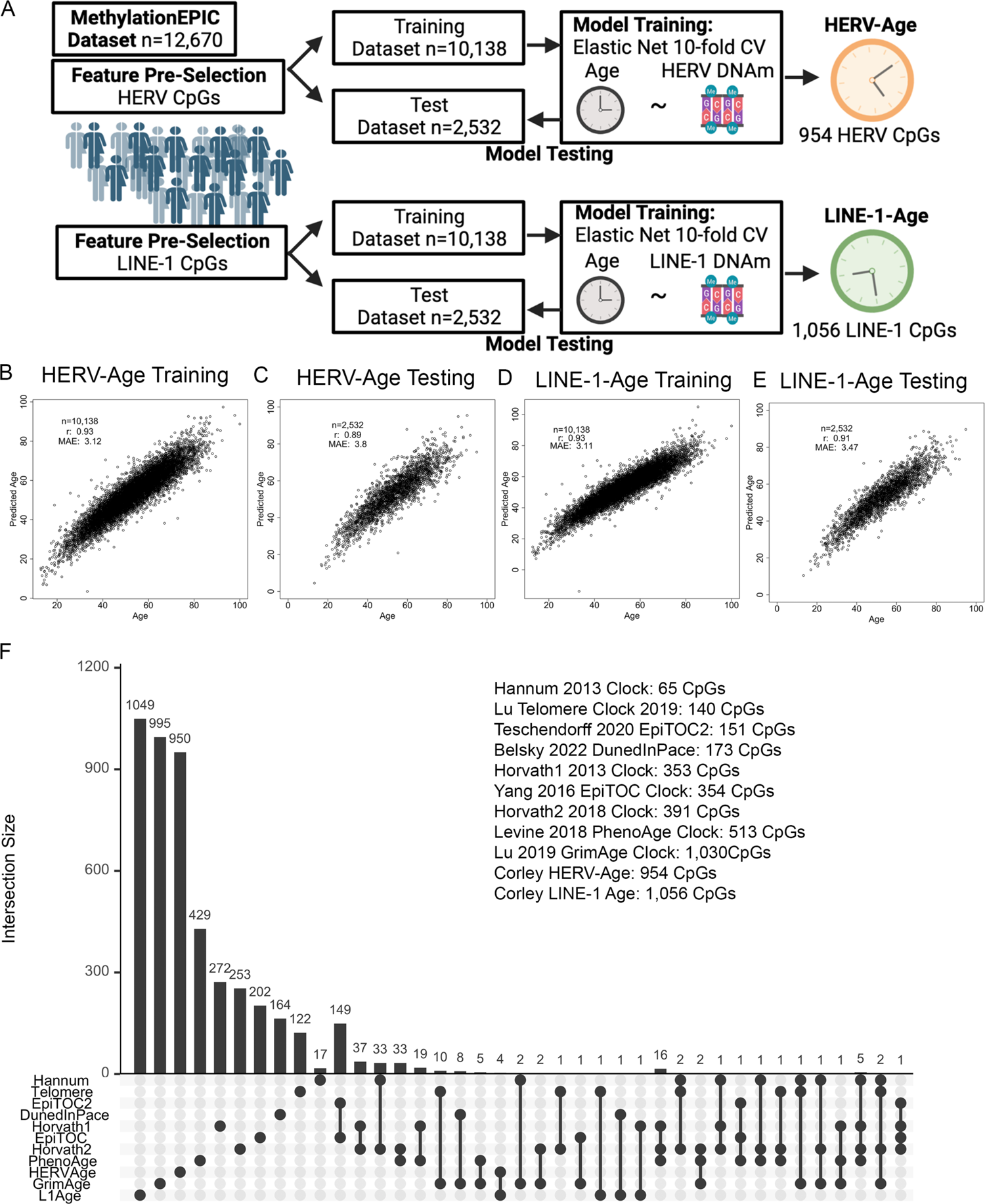
Construction of HERV-Age and LINE-1-Age Epigenetic Clocks. **A.** Diagram of workflow utilized to construct HERV-Age and LINE-1-Age. **B**. Age estimation 10 fold cross validation in training and **C.** test datasets for HERV-Age and **D-E.** LINE-1-Age. Panels report the sample size (n), the median absolute error (MAE), and Pearson correlation coefficient (r). **F**. Intersection plot of published epigenetic clocks CpGs, HERV-Age, and LINE-1-Age.

Next, we sought to examine whether the DNA methylation states of LINEs could also be utilized to generate an epigenetic clock of aging. Utilizing a similar elastic net regression approach with 10-fold cross-validation that we applied to HERV CpGs, we input LINE CpGs and preprocessed by splitting into an 80% training and 20% test data sets (**Fig. 2A**). This approach yielded a new LINE-based epigenetic clock (LINE-1-Age) based on 1,056 CpG sites (**Table S2**) located within LINEs that performed well in both training (**Fig. 2D**) and testing datasets (**Fig. 2E**) correlating significantly (r=0.93 training and r=0.91 testing) with chronological age. The MAE between LINE-1-Age and chronological age was 3.11 years in training and 3.47 years in testing datasets. In LINE-1-Age, we observed 8 CpGs each located at the L1FLnl_10q26.3e (cg03230038, cg09292354, cg09699225, cg13754569, cg17005927, cg18635896, cg20654619, and cg25120459) and L1FLnl_1p31.3z (cg00318766, cg08413469, cg08825216, cg16166651, cg18167921, cg19634693, cg20322613, cg26064634) LINEs. CpGs in L1FLnl_10q26.3e were downstream of the 3’UTR of the *ADGRA1* gene that encodes a G-protein-coupled receptor that has been shown to play an essential role for human pluripotent stem cells and reprogramming towards pluripotency(*36*). CpGs in L1FLnl_1p31.3z were located in an enhancer region of the *DEPDC1* gene that has been shown as a novel cell cycle related gene that regulates mitotic progression and has been shown to inhibit aging-related transcription factor *FOXO3*(*37*, *38*). L1FLnI_7q22.2a contained 5 CpGs in an enhancer region of the *ORC5* gene that is essential for the initiation of DNA replication and belongs to a family of origin recognition complex subunits that when depleted delay aging in budding yeast models(*39*). L1FLnI_18q23l contained 4 CpGs in the *NFATC1* gene that regulates activation, proliferation, differentiation, and programmed death of T cells(*40*). L1FLnI_Xp22.2h contained 4 CpGs that were in a enhancer and active transcriptional start region of the *TLR7* gene, part of a family of Toll-like receptors that show age-associated dysregulation of toll-like receptor signaling likely contributing to increased morbidity and mortality from infectious diseases in older people(*41*).

### HERV-Age and LINE-1-Age are constructed from unique CpGs when compared to existing epigenetic clocks

When building an epigenetic clock utilizing machine learning algorithms of large DNA methylation datasets, it is not typically known a priori which CpG sites are most informative for predicting age or other outcomes of interest(*11*, *42*). Hence, deconstructing epigenetic clocks and understanding the underlying biological mechanisms of these clocks has emerged as a major focus(*43*). To test whether retroelement CpGs could be utilized to construct an epigenetic clock, we relied on only utilizing manually annotated HERV and LINE-1 CpGs in the human genome. We sought to compare whether any of the 955 HERV-Age and 1,056 LINE-1-Age CpGs we identified overlapped with CpGs utilized in the construction of 9 existing epigenetic clocks that did not filter based on retroelement CpGs. We examined the intersection of HERV-Age and LINE-1-Age with Horvath’s multi-tissue predictor DNAmAge based on 353 CpG sites(*9*), the Horvath skin-and-blood clock based on 391 CpG sites(*44*), Levine’s DNAmPhenoAge based on 513 CpG sites(*16*), Hannum’s clock based on 71 CpG sites(*10*), the Lu’s telomere length predictor based on 140 CpGs(*45*), DNA methylation based mortality risk assessment GrimAge based on 1,030 CpGs(*15*), DunedinPace of aging based on 173 CpGs(*14*), and EpiTOC/EpiTOC2 mitotic clocks based on 354 and 151 CpGs(*46*). We found that the CpGs utilized in HERV-Age were all unique and did not overlap with any existing epigenetic clocks, suggesting that our HERV-Age clock captures novel biological DNA methylation features of aging not previously recognized (**Fig. 2F**). Notably, shared CpGs were observed among Horvath’s first-generation, second-generation clocks, and DunedInPace of aging suggesting the construction of these clocks captured some similar DNA methylation features of aging (**Fig. 2F**). Moreover, we observed that greater than 99% of CpGs in our new LINE-1-Age clock were also unique and did not overlap significantly with preexisting epigenetic clocks (**Fig. 2F**). LINE-1-Age shared 1 CpG with Lu’s Telomere clock, DunedInPace, and Horvath’s original epigenetic clock. Therefore, our retroelement-based epigenetic clocks capture a unique signal of human aging previously unconsidered by prior epigenetic clocks.

### Development of Composite Retroelement-Age Clocks

Since the DNA methylation states of both HERVs and LINEs could be utilized to construct highly predictive epigenetic clocks, we sought to develop a composite retroelement epigenetic clock (Retroelement-Age) by considering both the DNA methylation states of HERVs and LINEs. We utilized all the filtered CpGs based on our HERV and LINE annotations to construct a composite retroelement-based epigenetic clock (Retroelement-Age) based on a generalized linear elastic net model fit with 10-fold cross-validation on a 80% training dataset of 10,138 samples (**Fig. 3A**). The composite retroelement-age epigenetic clock consisted of 1,317 CpG sites (**Table S3**) and was highly predictive of chronological age in both training (r=0.95, MAE = 2.57) and test datasets (r=0.89, MAE = 3.81). Retroelement-Age consisted of multiple CpGs in regions belonging to the same retroelement including 10 CpGs located in the L1FLnl_8q22.2s element. Given the development of a new EPIC DNA methylation array(*47*), we sought to enhance the utility of a composite Retroelement-Clock for compatibility with both MethylationEPIC v.1.0 and MethylationEPIC v.2.0 data. Hence, we extended our manually curated annotation of Telescope-based MethylationEPIC CpGs to common probes on MethylationEPIC v2.0. We also included additional CpGs with LTR elements identified by RepeatMasker. Utilizing this annotation of compatible retroelement CpGs for MethylationEPIC v2.0, we constructed a new composite retroelement-based epigenetic clock (Retroelement-Age V2) based on a generalized linear elastic net model fit with 10-fold cross-validation on an 80% training dataset of 10,138 samples (**Fig. 3A**). The composite Retroelement-Age V2 epigenetic clock consisted of 1,378 CpG sites (**Table S4**) and was the most predictive of chronological age (**Fig. 3D-E**) compared to our prior retroelement-based clocks in both training (r=0.97, MAE = 1.87) and test datasets (r=0.96, MAE = 2.08). This was likely due to MethylationEPIC v2.0 containing more reliable probes(*47*, *48*). Our composite Retroelement-Age had 426 CpGs that overlapped with HERV-Age and 573 that overlapped with LINE-1 age. Notably, Retroelement-Age V2 was constructed from CpGs that did not overlap with existing first and second-generation epigenetic clocks (**Fig. 3F**).

**Fig. 3.**
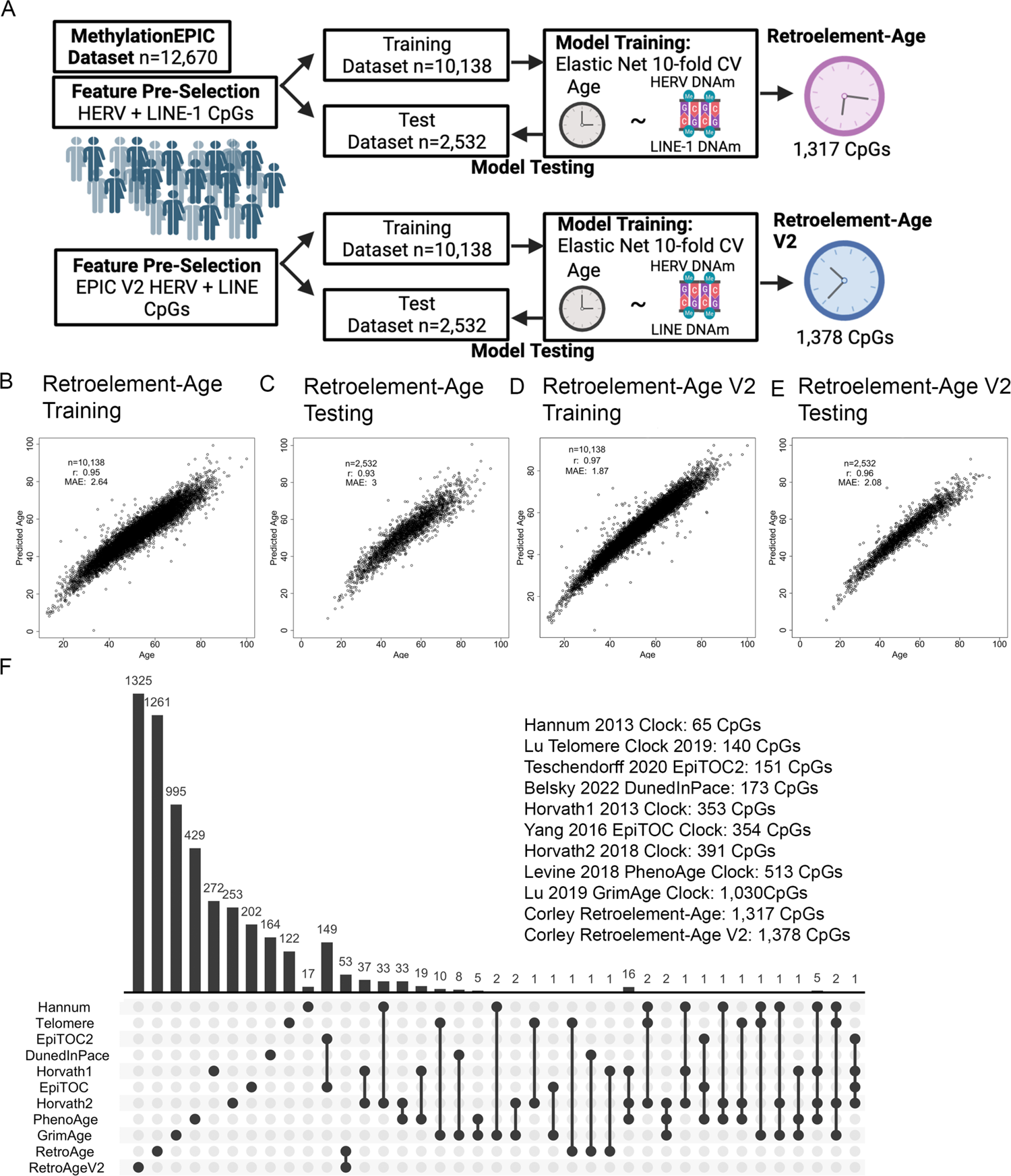
Construction of Composite Retroelement-Age and Retroelement-Age V2 Epigenetic Clocks. **A**. Diagram of workflow utilized to construct Retroelement-Age and Retroelement-Age V2. **B**. Age estimation 10 fold cross validation in training and **C.** test datasets for Retroelement-Age and **D-E.** Retroelement-Age V2 clocks. Panels report the sample size (n), the median absolute error (MAE), and Pearson correlation coefficient (r). **F**. Intersection plot of published epigenetic clocks CpGs, Retroelement-Age, and Retroelement-Age V2.

### Retroelement clocks are enriched in heterochromatin and quiescent chromatin regions containing H3K9me3, H3K9me2, and H3K4me1

We examined the overlap of CpGs in our retroelement clocks in specific regions of the epigenome based on consensus TFBSs, chromatin states, and consensus histone modifications. Consensus ChromHMM derived from 833 ENCODE ChromHMM calls from ENCODE version 2 was used(*24*). We found that CpGs in HERV-Age and LINE-1 Age clocks were enriched in inactive states consisting of constitutive heterochromatin, quiescent states, and zinc finger protein genes (ZNF/Rpts) (**Fig. 4A-F**). HERV-Age and LINE-1-Age were enriched in H3K9me3 and ATF7IP TFBSs (**Fig. 4A-F**). H3K9me3 deposition has been shown to be mediated by ATF7IP and SETDB1, which also are required for silencing retroelements(*49*). Retroelement-Age was enriched in KRAB zinc finger protein ZNF654, ATF7IP, ZBTB2, MAFG, quiescent/ heterochromatin states, and regions containing H3K9me2 and H3K9me3 (**Fig. 4G-I**). Collectively, identified TFBSs are supported by evidence in previous literature that demonstrates ZNFs to have co-evolved with retroelements to repress their activity(*50*). ZBTB2-binding dynamics *in vivo* are sensitive to differential DNA methylation and has been shown to repress the retrovirus HIV-1(*51*, *52*). and MAFG as a bidirectional regulator of transcription(*53*). Retroelement-Age V2 CpGs were enriched in MAFG and interaction partner NFE2L1(*54*), ZNF317, and ZNF654 (**Fig. 4J**). ZNF317 has been recently suggested to primarily target eutheria-specific ERVs(*55*). In addition to being enriched in quiescent chromatin states, Retroelement-Age V2 was enriched in active transcription start site (TSS)-proximal promoter states, enhancer states, and repressed Polycomb states (**Fig. 4K**). Polycomb group proteins are hypothesized as an evolutionarily conserved mechanism to silence transposable elements(*56*). Retroelement-Age V2 CpGs were enriched in H3K4me1 regions, which is both repressive and marker of poised and active enhancers(*57*, *58*).

**Fig. 4.**
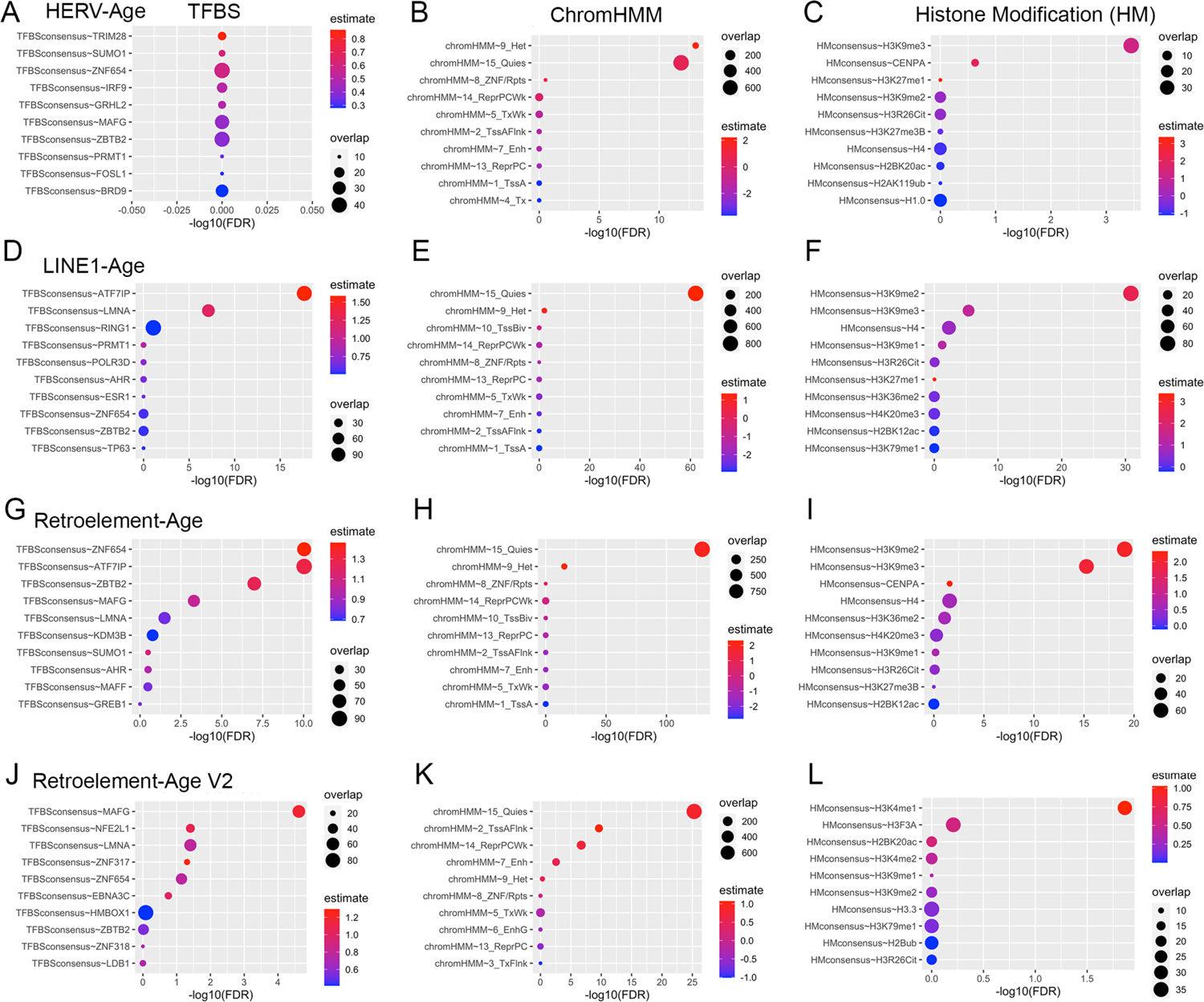
Enrichment plot of retroelement-based epigenetic clocks in consensus transcription factor binding sites (TFBS), ENCODE ChromHMM chromatin states, and histone modifications for **A-C**. HERV-Age, **D-F**. LINE-1-Age, **G-I**. Retroelement-Age, and **J-L.** Retroelement-Age V2. Fisher’s exact test, estimate represents fold enrichment, and overlap the number of CpGs.

### Associations of retroelement-based clocks to existing epigenetic clocks and DNA methylation inferred immune cell compositions

We utilized the DNA methylation dataset from 12,670 participants with a known chronological age to examine the relationships between chronological age, our new retroelement clocks (HERV-Age, LINE-1-Age, Retroelement-Age, Retroelement-Age V2), and existing epigenetic clocks. Despite being constructed from largely unique CpGs compared to previous epigenetic clocks, we found that HERV-Age (r=0.92), LINE-1-Age (r=0.94), Retroelement-Age (r=0.94), and Retroelement-Age V2 (r=0.97) significantly associated with chronological age to a similar or better degree than existing epigenetic clock algorithms such as Horvath1 (r=0.89), Hannum (r=0.91), and PhenoAge (r=0.85) (**Fig. 5A**). We also observed this relationship was similar or better than new principal component-based versions of epigenetic clocks(*59*) including PCHorvath1 (r=0.88), PCHorvath2 (r=0.84), PCHannum (r=0.90), PCPhenoAge (r=0.88), and PCGrimAge (r=0.96) (**Fig. 5A**). Next, we utilized an enhanced 12 leukocyte cell deconvolution algorithm to infer the immune cell proportions in all 12,670 participants(*60*). We found that increasing HERV-Age, LINE-1-Age, Retroelement-Age, and Retroelement-Age V2 significantly associated with a decreased inferred proportion of naive CD8+ T cells (r=-(0.45-0.46)) and CD4+ T cells (r=-(0.16-.018)) (**Fig. 5B**). These findings underscore the potential of our novel retroelement-based epigenetic clocks as novel indicators for aging and age-related immune system dynamics.

**Fig. 5.**
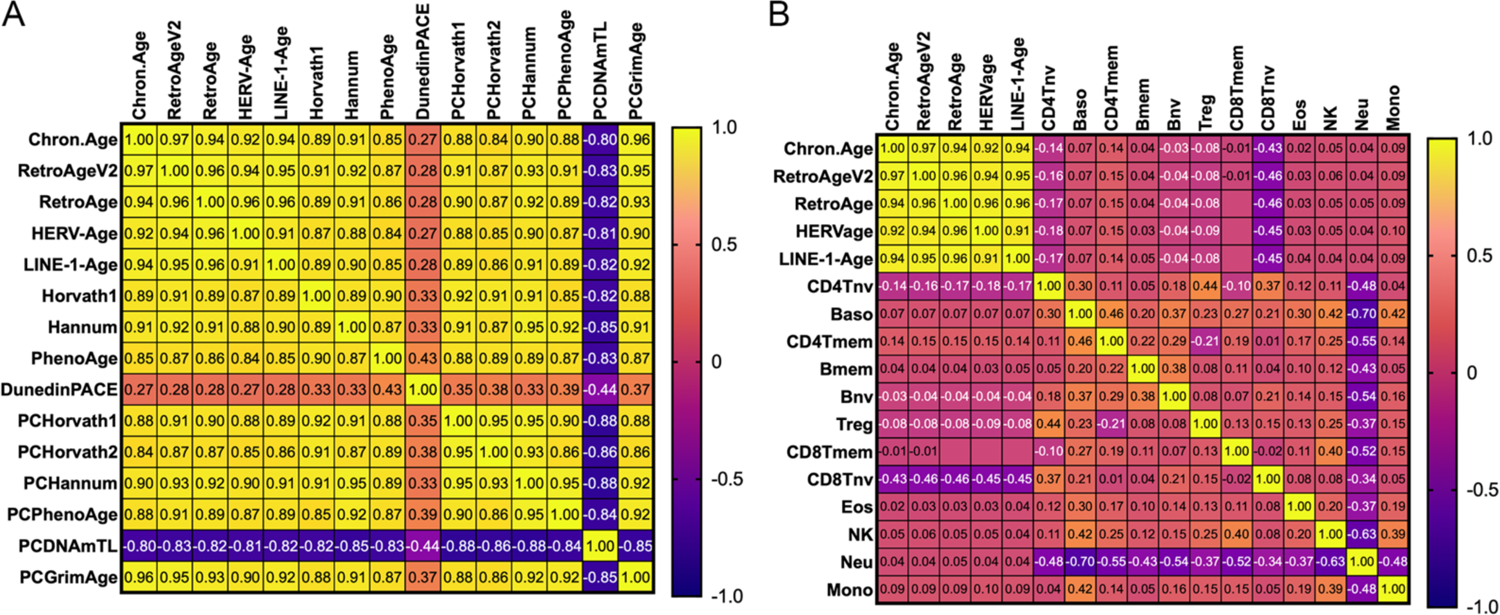
Correlograms of chronological age of 12,670 people, A. epigenetic clock age estimates, and B. inferred immune cell subsets.

### Independent validation and reliability of retroelement-based epigenetic clocks

We sought to evaluate the performance and validate our 4 retroelement-based epigenetic clocks in a completely independent and external DNA methylation dataset. We obtained a blood MethylationEPIC v1.0 dataset from 790 individuals spanning chronological ages of 55-95 years (55.94% male) in the ADNI cohort(*61*). In this dataset, Retroelement-Age V2 (r=0.83), Retroelement-Age (r=0.72), LINE-1-Age (r=0.70), and HERV-Age (r=0.62) correlated with chronological age (**Fig. 6A-D**). We evaluated test-retest reliability of RepeatAge V2 using 30 replicate blood-sample MethylationEPIC v1.0 samples based on the intraclass correlation coefficient (ICC) using a linear mixed-effects model. We found that reliability was high with an ICC of 0.996 (**Fig. 7A**).

**Fig. 6.**
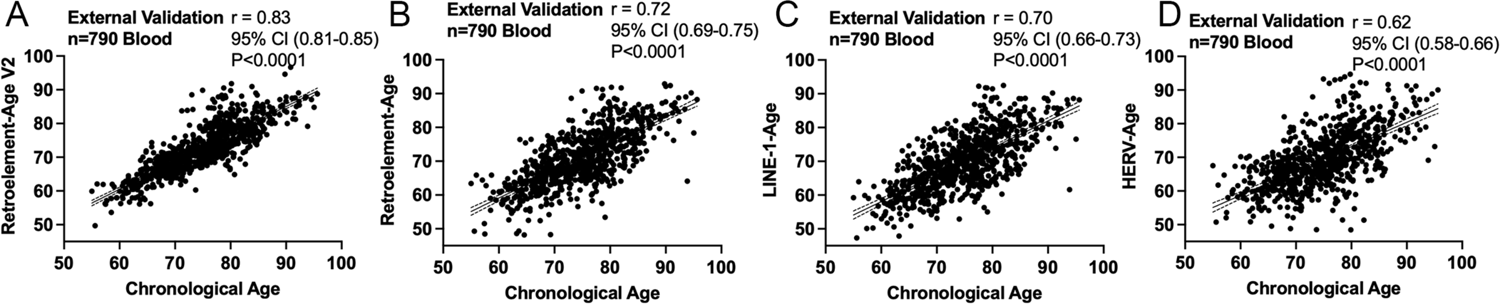
External validation of retroelement-based epigenetic clocks. Scatterplots of **A.** Retroelement-Age V2, **B.** Retroelement-Age, **C.** LINE-1-Age, and **D.** HERV-Age and chronological age in 790 people. Independent data generated by the Alzheimer’s Disease Neuroimaging Initiative study.

**Fig. 7.**
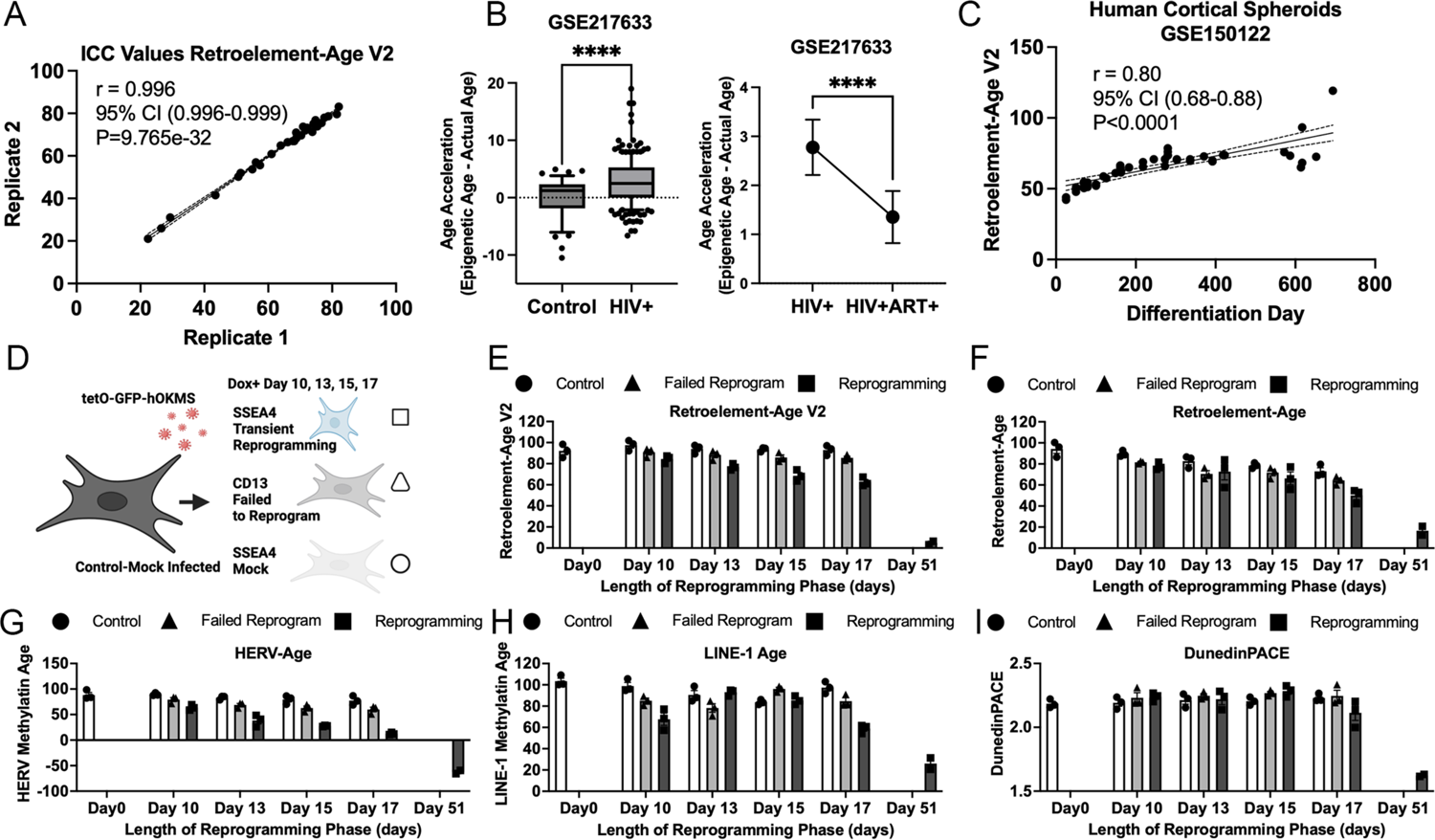
Reliability and applications of retroelement-based epigenetic clocks. **A.** Intra-class correlation coefficient (ICC) of Retroelement-Age V2 on 30 replicate blood sample MethylationEPIC v1.0 samples. **B.** Age acceleration detected utilizing Retroelement-Age V2 in external data from people living with HIV compared to healthy controls. Retroelement-Age V2 responsive to antiretroviral therapy (ART) in people living with HIV followed longitudinally before and after 96 weeks of ART. **C.** Application of Retroelement-Age V2 in tracking human cortical brain organoids culture age. **D.** Application of retroelement-based epigenetic clocks to transient reprogramming-induced rejuvenation strategies dataset GSE165179. Epigenetic age estimates for Control (circle symbol), failed to program (triangle symbol), and transiently reprogrammed (square symbol) fibroblasts at Day 0, 10, 13, 15, and 17 based on **E.** Retroelement-Age V2, **F.** Retroelement-Age, **G.** HERV-Age, **H.** LINE-1-Age, and **I** DunedInPace.

### Effect of HIV-1 infection and antiretroviral therapy on retroelement-based epigenetic clocks

Prior studies have reported accelerated epigenetic aging in people living with infectious diseases such as HIV-1(*62*, *63*). We tested whether our retroelement clocks would capture accelerated epigenetic aging in a MethylationEPIC dataset from a cohort of 185 people living with HIV-1 (with pre-ART and post-ART longitudinal samples) and 44 demographically matched healthy controls (GSE217633)(*64*). We replicated evidence of epigenetic age acceleration in people living with HIV-1 pre-ART. We found a significant epigenetic age acceleration of mean average 2.8 years in people living with HIV-1 compared to epigenetic age difference from chronological age of −0.19 years in healthy controls (**Fig. 7B**). Prior work has shown several FDA approved antiretroviral drugs can effectively inhibit HERV-K(*65*). Hence, we hypothesized that antiretroviral therapy utilized to treat HIV-1 would reduce retroelement-based epigenetic age. We found a significant reduction in epigenetic age following 96 weeks of antiretroviral therapy comparing longitudinal samples of people living with HIV-1 at pre- and post-ART timepoints (**Fig. 7B**).

### Application of Retroelement-based epigenetic clocks to human cortical organoids during long term maturation

Prior work has shown the utility in epigenetic clocks in predicting culture age of human cortical organoids maintained in long-term cultures up to 694 days(*66*). We applied Retroelement-Age V2 to Methylation DNA methylation data (GSE150122) across 13 timepoints generated for six human induced pluripotent stem cell (hiPSC) lines obtained from five individuals (four male and one female) that were differentiated to human cortical spherioids (hCS)(*67*). We observed a significant relationship (r= 0.80, P<0.0001) between the *in vitro* culture age of hCS and predicted Retroelement-Age V2 (**Fig. 7C**). These findings are consistent with application of Horvath’s epigenetic clock(*66*) and suggest a dynamic relationship between DNA methylation states of human endogenous retroviruses and induced pluripotent stem cells(*68*).

### Transient reprogramming-induced rejuvenation strategies utilizing Yamanaka factors reverse Retroelement-based epigenetic clocks

Transient reprogramming has emerged as a controversial strategy to epigenetically rejuvenate cells as an anti-aging strategy(*43*, *69*). A key outcome measures utilized to assess epigenetic rejuvenation through transient reprogramming has been epigenetic clocks. However, findings suggests some clocks may not have utility in assessing epigenetic rejuvenation achieved by transient reprogramming(*43*, *69*). Based on prior work showing HERVs are a key mechanism involved in involved in human iPSC generation and re-establishment of differentiation potential of induced pluripotent stem cells (iPSCs)(*68*), we sought to examine whether our retroelement-based epigenetic clocks would inform transient reprogramming-induced rejuvenation strategies. We leveraged a published epigenetic rejuvenation DNA methylation dataset(*69*) GSE165179 and calculated epigenetic ages for control, failed to program, and transiently reprogrammed fibroblasts at Day 0, 10, 13, 15, and 17 (**Fig. 7D**). Retroelement-Age V2, Retroelement-Age, HERV-Age, and LINE-1-Age were significantly reversed in reprogrammed cells compared to control cells at Day 10, 13, 15, and 17 (**Fig. 7E-F**). We observed that the DunedinPACE epigenetic clock(*14*) was not that responsive to assessing transient reprogramming (**Fig. 7I**). Application of PC-based epigenetic clocks showed PCHorvath1, PCHorvath2, and PCHannum detected transient reprogramming at days 13, 15, and 17. More reliable mortality estimate clocks PCPhenoAge and PCGrimAge were less responsive to transient reprogramming suggesting decreased utility in being used to assess the success of transient reprogramming. These findings suggest retroelement-based epigenetic clocks as an additional outcome measure in ongoing transient reprogramming efforts and externally validate prior findings showing the utility of some epigenetic clocks to assess epigenetic reprogramming and anti-aging strategies(*69*). Whether a new transient-native-treatment (TNT) reprogramming strategy shown to correct aberrant transposable element expression impacts retroelement-based epigenetic clocks remains unclear(*70*).

### Immune Transcriptome Telescope-Age

We sought to evaluate whether age-related RNA expression of locus-specific retroelements in the immune system overlapped with DNA methylation loci identified in our retroelement clocks. Transcriptomic age signatures have been shown to offer a complementary predictor of biological aging(*71*). We utilized the Telescope computational pipeline(*18*) to estimate HERV and LINE element expression resolved to specific genomic locations in an independent RNA-Seq dataset of blood from 157 human donors spanning chronological ages 20-74 year (GEO: GSE193141) (**Fig. 8A**). Using a generalized linear elastic net model fit with 10-fold cross-validation on an 80% training and 20% test dataset using both HERV and LINEs expression in blood, we found that the expression levels of 95 retroelements (**Table S5**) were highly predictive of chronological age (r=0.98 training and r=0.62 testing) (**Fig. 8B-C**). Among the elements, we observed 41 LINEs, supporting hypotheses regarding the resurrection of particular retroelement in the aging process(*6–8*). Integrating loci from our HERV-Age and LINE-1-Age clocks, we found overlap of age-associated DNA methylation and RNA changes in 11 locus specific HERV and LINE-1 elements including ERVLB4_11p15.5b, ERVLE_1q25.3d, HERV3_19p13.3, HERV4_5q22.1, HERVFRD_2p12a, HERVIP10FH_11p15.5, L1FLnI_12q13.3b, L1FLnI_1q31.3h, L1FLnI_2q37.2b, L1FLnI_8q22.3c, and MER4B_14q32.33d (**Fig. 8D**). These findings suggest that a subset of the loci identified in retroelement-based epigenetic clocks may relate to epigenetic dysregulation leading age-related transcriptional dysfunction of specific retroelements.

**Fig. 8.**
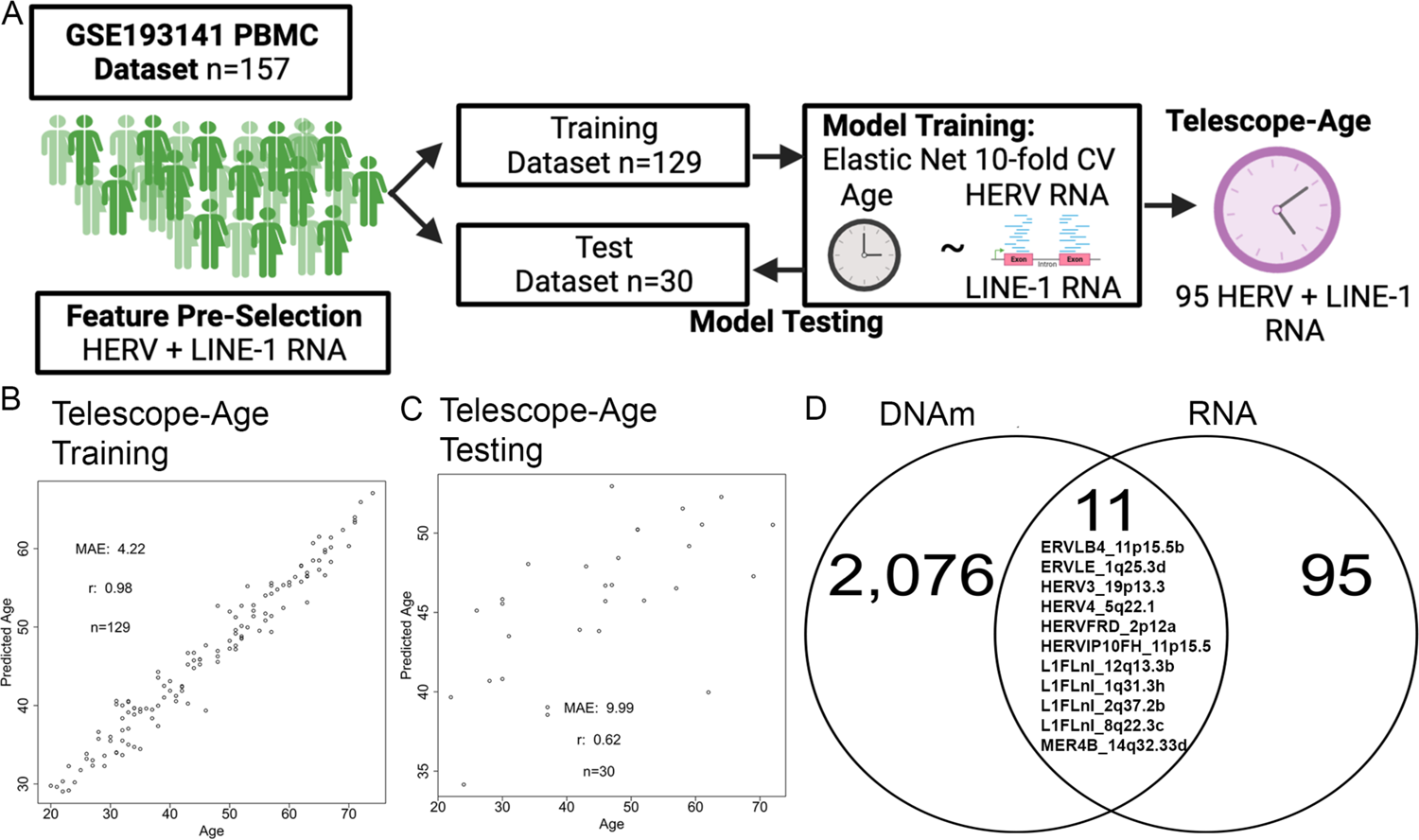
Integration of Age-related transposable element expression and retroelement-based DNA methylation epigenetic clocks. **A.** Diagram of workflow utilized to construct immune transcriptome Telescope-Age. **B**. Age estimation 10-fold cross validation in training and **C.** test datasets for Telescope-Age. **D.** Overlap of HERVs and LINEs identified in retroelement-based DNA methylation epigenetic clocks and transcriptome immune system datasets.

### Evidence of Human Multi-Tissue Retroelement clock

We sought to examine whether DNA methylation states of retroelements could be utilized beyond the immune system to capture human aging. To test this hypothesis, we used publicly available Genotype-Tissue Expression (GTEx) DNA methylation data for 987 human samples from nine tissue types spanning breast mammary tissue, muscle skeletal, lung, ovary, kidney, testis, prostate, colon, and whole blood(*19*). To test whether there was evidence of a human multi-tissue retroelement clock, we utilized all the filtered CpGs based on our HERV and LINE-1 annotations and constructed a new composite retroelement-based epigenetic clock (Retroelement-TissueAge) based on a generalized linear elastic net model fit with 10-fold cross-validation on an 80% training and 20% validation dataset. Since this DNA methylation dataset consisted of multiple tissues from the same donor, we sought to minimize data leakage and ensured each donor was only in one of the sets (either training or validation). The Multi-Tissue Retroelement epigenetic clock consisted of 734 CpG sites (**Table S6**) and was highly predictive of chronological age in both training (r=0.99, MAE = 0.76) and test datasets (r=0.75, MAE = 5.07) (**Fig. 9A-B**).

**Fig. 9.**
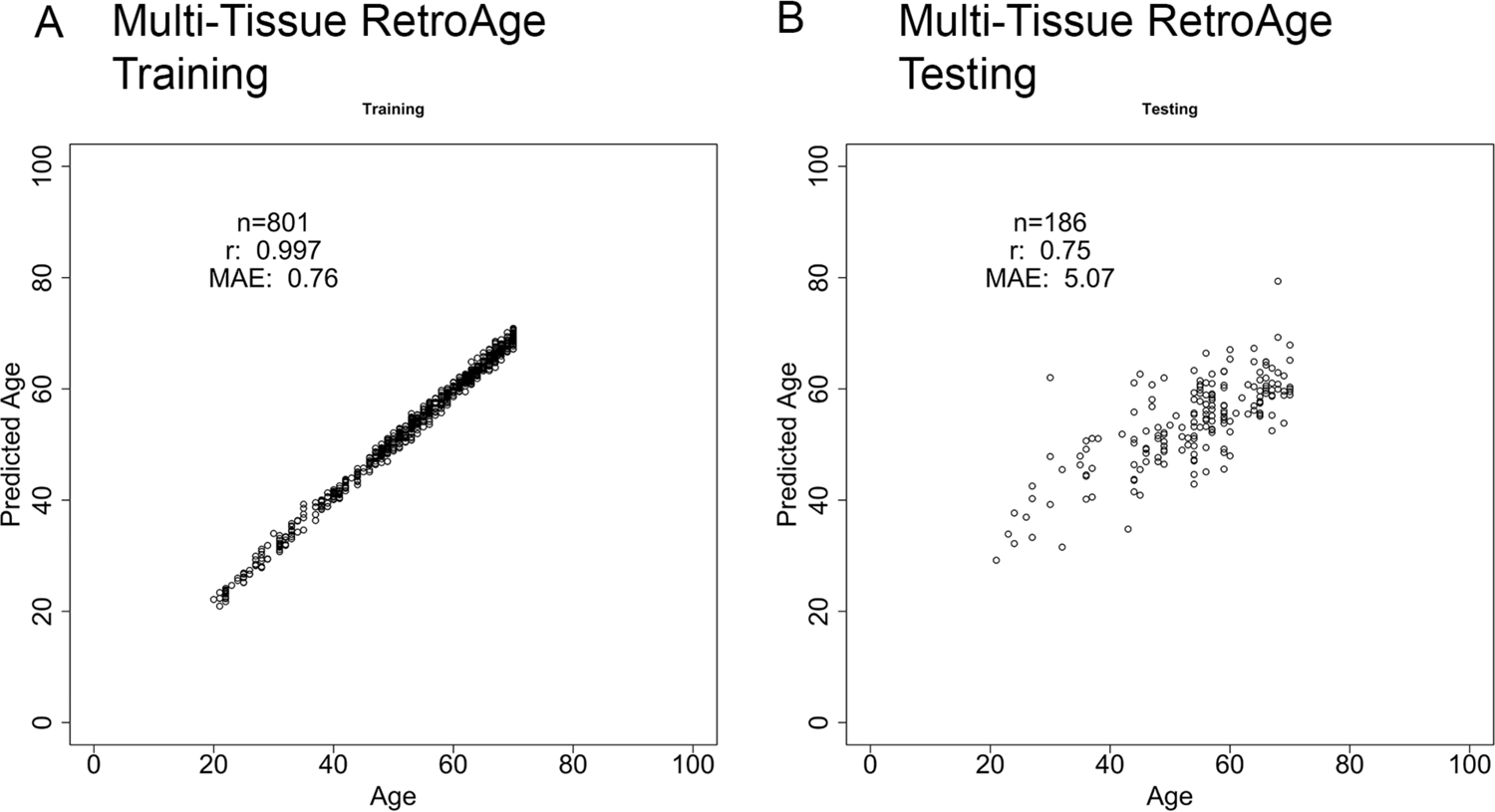
Integration of Age-related transposable element expression and retroelement-based DNA methylation epigenetic clocks. **A**. Age estimation 10 fold cross validation in training and **B.** test datasets for Multi-Tissue RetroAge GTEx project data GSE213478.

### Evidence of Pan-Mammalian Species Multi-Tissue Retroelement Clock

Recent data from the international Mammalian Methylation consortium demonstrated the development of universal pan-mammalian clocks using DNA methylation data from over 11,000 samples spanning 59 tissue types and across 185 mammalian species using the HorvathMammalMethylChip40 (Mammal40) array, a mammalian array that targets over 36,000 conserved CpGs across mammalian species(*20*, *72*). However, the specific CpGs utilized to construct these pan-mammalian clocks did not focus on CpGs located within retroelements or other transposable elements. Hence, we first sought to evaluate whether the Mammal40 platform contained probes overlapping with transposable elements. We used RepeatMasker annotated probes in the Mammal40 platform and identified 563 loci in the class of DNA, LINE, LTR, SINE, and Unknown. Next, we utilized the pan-mammalian DNA methylation dataset consisting of 12,146 samples and relative ages calculated as log transformed chronological ages with an offset of 2 years added. First, we utilized all 563 DNA, LINE, LTR, SINE, and unknown CpGs identified by RepeatMasker annotated probes and developed a pan-mammalian species clock consisting of 398 CpG sites (**Table S7**) that was highly predictive of chronological age in both training (r=0.94) and test datasets (r=0.94) (**Fig. 10A-B**). Next, we only used the 211 CpGs in LINEs and LTR and found we could also construct a highly predictive pan-mammalian species clock based on only 201 CpGs in HERVs and LINEs (**Table S8**) covered in the dataset (**Fig.10C-D**). We examined the intersection of the CpGs utilized in our pan-mammalian TransAge and RetroAge clocks with the published 3 universal pan-mammalian clocks(*20*). Notably, we found no overlap with CpGs utilized to construct the currently available pan-mammalian epigenetic clocks (**Fig. 10E**), suggesting additional insights into the evolutionary components of aging-associated processes not previously detected.

**Fig. 10.**
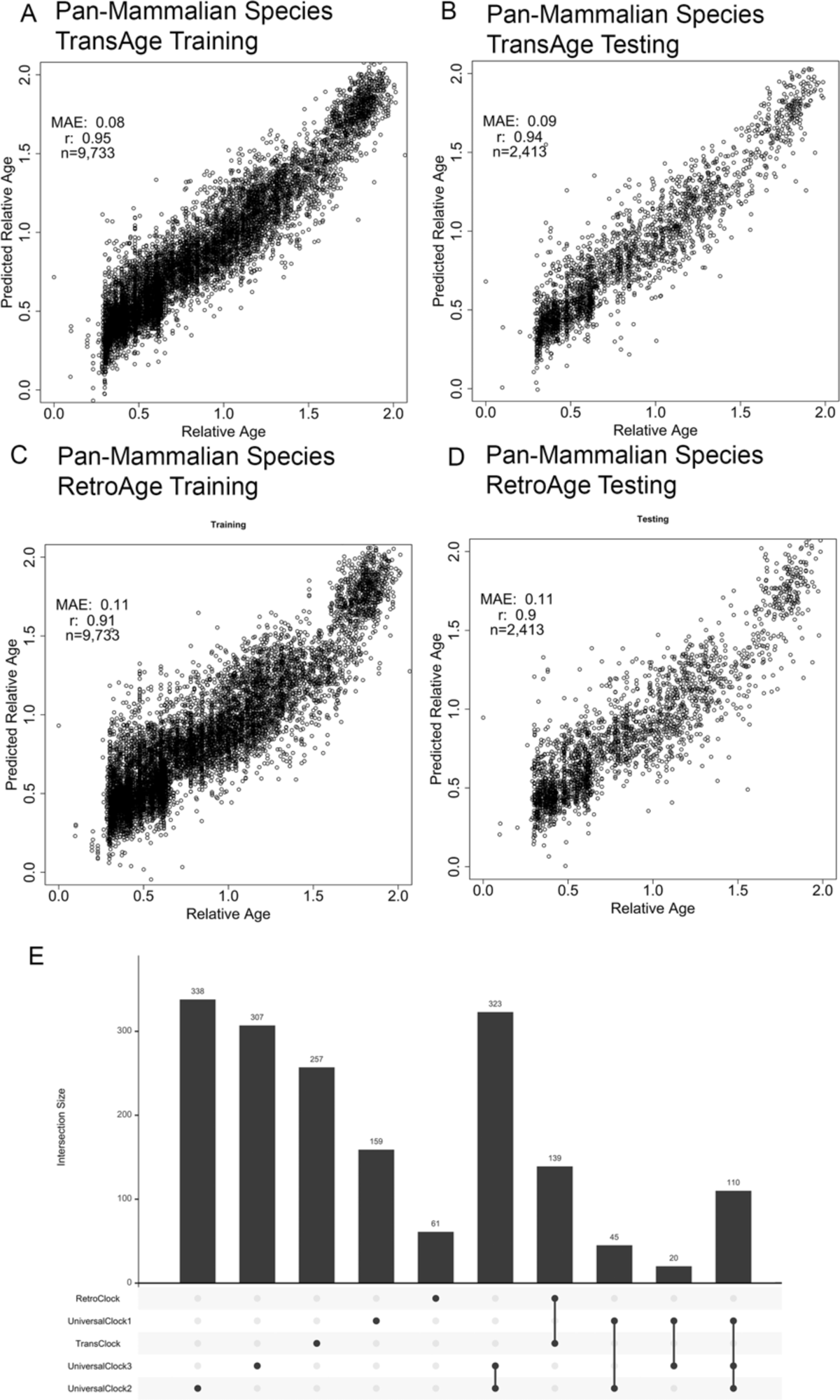
Construction of Pan-Mammalian Transposable element and Retroelement-based Epigenetic Clocks. **A**. Age estimation 10 fold cross validation in training and **B.** test datasets for TransAge and **C-D.** RetroAge clocks. Panels report the sample size (n), the median absolute error (MAE), and Pearson correlation coefficient (r). **E**. Intersection plot of published Pan-mammalian epigenetic clocks CpGs, Pan-Mammalian TransAge, and Pan Mammalian RetroAge.

## Discussion

Distinct DNA methylation patterns in the human genome can reliably estimate a person’s biological age, thereby leading to the development of a myriad of epigenetic clocks that have emerged as promising biomarkers of biological age for the geroscience field. While epigenetic age has been linked to specific hallmarks of aging(*73*), the role of epigenetic clocks in mechanisms of aging linked to retroelements remains unclear. Here, we show the potential of DNA methylation states in HERVs and LINEs as highly accurate epigenetic clocks of immune system aging, supporting the hypothesis that interspersed repeat elements may be involved in aging. Using only HERV and LINE-1 DNA methylation states, we developed highly accurate retroelement-based epigenetic clocks that minimally overlapped with preexisting first and second-generation epigenetic clocks. We find evidence of clock CpGs used in our retroelement-based epigenetic clocks located in HERV elements previously linked to aging and immune function, such as HERV-K and MER41. We also captured retroelement-based epigenetic clock CpGs in novel retroelements not previously linked to aging, suggesting a broader role of locus-specific retroelements in biological aging. Applying these retroelement-based epigenetic clocks to DNA methylation datasets, we find that retroelement clocks are reversed during transient epigenetic reprogramming(*69*), accelerated in people living with HIV(*74*), responsive to antiretroviral therapy(*74*), and accurate in estimating long-term culture ages of human brain organoids(*66*). Finally, we demonstrate that retroelement clocks extend to diverse human tissues and across mammalian species using DNA methylation from GTEx(*19*) and from the Mammalian Methylation Consortium(*20*). Together, these findings support the hypothesis of dysregulation of endogenous retroelements as a potential contributor to the biological hallmarks of aging and suggest that therapeutic interventions modifying the epigenetic states of specific retroelements in the human genome could have beneficial effects against a root cause of aging. Additionally, these studies suggest that retroelement-based epigenetic clocks are evolutionarily conserved throughout mammals, and that further study of which throughout post-speciation events could provide valuable insight into the evolutionary components of aging-associated processes and the impact of paleovirology on each species’ lifespan.

Emerging evidence suggests a correlation between aging and the reactivation of specific retroelements, primarily LINEs and HERV-K-derived retrovirus like particles (RVLPs). Prior work has shown that the transcriptional derepression of LINEs in immune cells cells triggers interferon production to contribute to inflammaging (*7*, *75–77*). Recent findings have suggested a key role of the endogenous retrovirus HERV-K (HML-2) activation in cellular senescence and tissue aging(*6*). Data from Drosophila models of aging have revealed that stimulating retrotransposon activity increases mortality and accelerates a subset of aging phenotypes(*78*). Our data extend these findings and suggest DNA methylation changes of locus-specific retroelements plays a key role in immune aging and potentially eliciting biological hallmarks of aging.

A notable observation was that the set of CpGs that were utilized to construct our retroelement-based epigenetic clocks did not show a significant overlap with pre-existing first and second-generation epigenetic clock algorithms(*9*, *14–16*). Differences between clocks have arisen from the focus on DNA methylation data compatibility, considerations of probe reliability, datasets utilized, and being trained to predict different aging-related variables, such as chronological age, composite biomarkers of aging, mortality risk, or mitotic divisions. Our findings suggest that retroelement-based epigenetic clocks capture previously undetected facets of biological aging to complement current epigenetic clocks that appear to capture distinct aspects of aging and associate with different biological hallmarks of aging, environmental exposures, traits, and disease patterns (*11*, *79–81*).

Existing iterations of epigenetic clocks were constructed based on a hypothesis about dynamic DNA methylation changes being associated with age, health biomarkers, and mortality. The filtering of CpGs utilized to construct prior epigenetic clocks was typically performed to account for reliable probes and Illumina methylation array platform compatibility. Moroever, we found that using a limited set of CpGs from the HorvathMammalMethylChip40 (Mammal40) array that only contained 643 loci based on RepeatMasker in the class of DNA, LINE, LTR, SINE, and Unknown we were able to construct a highly accurate Pan-mammalian retroelement methylation clock based on 220 CpGs. Less than 3% (6 CpGs) overlapped with CpGs utilized to construct the 3 Universal Pan-Mammalian methylation clocks by Lu et al.(*20*). These additional findings suggest an evolutionarily conserved dysregulation of DNA methylation states within retroelements may play a role beyond mammalian development into aging(*82*).

We developed a new composite retroelement-based epigenetic clock (Retroelement-Age V2) that contains CpGs covered on Illumina’s MethylationEPIC v2.0 kit. An interesting feature of Retroelement-Age V2 is that none of the CpGs utilized to construct this clock overlapped with 9 existing first and second-generation epigenetic clocks. We found a significant enrichment of 80 CpGs in this clock at CCCTC-binding factor (CTCF) sites of the genome. These findings suggest that DNA methylation changes in retroelements, particularly at CTCF (CCCTC-binding factor) binding sites, may potentially play a role in aging. CTCF is a critical protein involved in the organization of chromatin structure and the regulation of gene expression. It acts as an insulator, helping to define boundaries between chromatin domains and regulate the accessibility of genes to the transcriptional machinery. Our findings support prior DNA methylation work showing changes with aging observed in CTCF binding sites(*83*) and a pan-tissue DNA methylation epigenetic clock based on deep learning that found that the most important CpG sites were proximal to CTCF binding sites(*84*). Future work will need to examine whether modifying DNA methylation states at retroelements overlapping with CTCF binding impacts immune aging by changes in transcriptional regulation, insulation of chromatin domains, and the organization of higher-order chromatin structure.

Prior epigenetic clocks detect accelerated aging effects related to infection with the exogenous retrovirus HIV-1 (*62*, *63*, *85–87*). People living with HIV-1 are a population who exhibit increased features of biological aging (*88–90*), likely due to infection from the exogenous retrovirus HIV-1, chronic inflammation, antiretroviral therapy, and lifestyle effects (*91–93*). Prior work has shown that HIV-1 infection activates certain HERVs including HERV-K. This activation may be an additional factor contributing to accelerated/attenuated aging in people living with HIV related to immune dysfunction, inflammation, and immunosenescence. Prior epigenetic clocks did not provide insights into whether altered DNA methylation states of HERVs or LINEs related to HIV-1 infection. By applying our retroelement-based epigenetic clocks, we find evidence that suggest an increase in the epigenetic age compared to chronological age related to HIV, supporting the hypothesis that retroviruses accelerates biological aging. Additionally, examining longitudinal data from people living with HIV-1 (PLWH) receiving antiretroviral therapy(*64*), we find that antiretroviral therapy treatment significantly reverses HIV-1-related increased retroelement-based epigenetic age. Whether antiretroviral therapy can be used as therapeutic to improve health and increase lifespan by reversing retroelement-based epigenetic age in the absence of exogenous retroviral infections therefore warrants further investigation. Analysis of the effects of pre-exposure prophylaxis (PrEP), which delivers antiretroviral drugs to reduce the risk of HIV infection, on retroelement-based epigenetic age can uncover potential effects of antiretroviral drugs in HIV-negative people on aging.

A limitation of our findings is that our study primarily focused on DNA methylation datasets from the human immune system. While we included DNA methylation data from GTEx samples across nine tissues, this dataset was limited and did not permit the construction of systems-specific retroelement-based epigenetic clocks. Our application of retroelement-based epigenetic clocks across mammalian species was also limited to highly conserved CpG DNA sequences and could benefit from a species-specific analysis that considers uniquely arisen CpG sites in respective species genomes.

In summary, these findings highlight the potential of DNA methylation states of specific retroelements as reliable predictors of biological aging, complementing existing epigenetic clocks and offering an additional mechanism to consider in epigenetic clock signals. Collectively, our results suggest a renewed emphasis on the role of retroelements in human aging and warrants further study on their undefined roles in geroscience.

## Materials and Methods

### Discovery cohort

The TruDiagnostic Biobank cohort, previously described in Chen et al(*94*), included 13,109 individuals who took the commercial TruDiagnostic TruAge test and had their DNA methylation data generated from whole blood. The participants were recruited between October 2020 and April 2023 and were predominantly from the United States.

### MethylationEPIC V1.0 DNA methylation pre-processing and analysis

Peripheral blood samples were collected using a lancet and capillary method and placed in a lysis buffer for DNA extraction. Then, 500 ng of DNA was treated with bisulfite using the EZ DNA Methylation kit from Zymo Research following the manufacturer’s instructions. The bisulfite-treated DNA samples were randomly assigned to a well on the Infinium HumanMethylationEPIC BeadChip, which was then amplified, hybridized, stained, washed, and imaged with the Illumina iScan SQ instrument to obtain raw image intensities. To pre-process the TruDiagnostic methylation data, we used the *minfi* pipeline(*95*), and low quality samples were identified using the *qcfilter()* function from the ENmix package(*96*), using default parameters. A total of 12,670 individuals, representing 96.7% of the original samples, passed the QA/QC (p < 0.05) and were deemed to be high quality samples.

### HERV, LINE-1, and Composite Retroelement Clocks Construction

CpGs covered on the Illumina Infininium MethylationEPIC (EPIC) V1 array were filtered using a manually curated locus-specific HERV annotation of 60 HERV families(*18*). This annotation approach identified that 5,436 probes (0.62% of EPIC) assessed DNA methylation at a CpG within a HERV. A locus-specific LINE-1 annotation of 13,545 loci derived from L1Base(*18*) was used to annotate probes included on the EPIC array. This annotation approach identified that 5,543 probes (0.64% of EPIC) assessed DNA methylation at a CpG within a LINE-1 element. The locus specific HERV and LINE-1 annotations were then used to filter two beta matrixes of CpGs for all 12,670 samples. The caret R package was used to load metadata for chronological age of all 12,670 samples. The HERV and LINE-1 filtered beta matrixes were then split into 80% training and 20% validation datasets. The glmnet R package was then used to train models for HERV and LINE-1 clocks with a 10-fold validation utilizing an elastic net. The UpSetR R package was used for visualization of intersection of CpGs(*97*, *98*).

### Multi-Tissue and Pan-Mammalian Species Retroelement Clocks Construction

Retroelement filtered beta matrixes were for a GTEx dataset of 987 samples (GSE213478) and Pan-Mammalian Species dataset of 12,146 samples (GSE223748) were split into 80% training and 20% validation datasets. Chronological ages and log transformed chronological ages with an offset of 2 years added were used. The glmnet R package was then used to train models for clocks with a 10-fold validation utilizing an elastic net.

### CpG Feature Enrichment

The knowYourCG tool was utilized for examining CpG feature enrichment using Illumina probe IDs and databases associated with certain CpGs available at https://github.com/zhou-lab/KYCG_knowledgebase_EPIC/tree/main/Studies. Fisher’s exact test was utilized to test CpG enrichments. Enrichment results were visualized using the KYCG_plotDot function in SeSAMe.

### Public datasets

Data used for validation and testing of retroelement clocks was publicly available via Gene Expression Omnibus (GEO). MethylationEPIC V1 array IDATs were downloaded from GEO using the following accession numbers: GSE165179, GSE150122, GSE213478, GSE223748. All GTEx protected data was accessed via the GTEx Portal. Validation DNA methylation data used in the preparation of this article were obtained from the Alzheimer’s Disease Neuroimaging Initiative (ADNI) database (adni.loni.usc.edu). As such, the investigators within the ADNI contributed to the design and implementation of ADNI and/or provided data but did not participate in the analysis or writing of this report. A complete listing of ADNI investigators can be found at http://adni.loni.usc.edu/wp-content/uploads/how_to_apply/ADNI_Acknowledgement_List.pdf. Data collection and sharing for this project was funded by the Alzheimer’s Disease Neuroimaging Initiative (ADNI) (National Institutes of Health Grant U01 AG024904) and DOD ADNI (Department of Defense award number W81XWH-12-2-0012). ADNI is funded by the National Institute on Aging, the National Institute of Biomedical Imaging and Bioengineering, and through generous contributions from the following: AbbVie, Alzheimer’s Association; Alzheimer’s Drug Discovery Foundation; Araclon Biotech; BioClinica, Inc.; Biogen; Bristol-Myers Squibb Company; CereSpir, Inc.; Eisai Inc.; Elan Pharmaceuticals, Inc.; Eli Lilly and Company; EuroImmun; F. Hoffmann-La Roche Ltd and its affiliated company Genentech, Inc.; Fujirebio; GE Healthcare; IXICO Ltd.; Janssen Alzheimer Immunotherapy Research & Development, LLC.; Johnson & Johnson Pharmaceutical Research & Development LLC.; Lumosity; Lundbeck; Merck & Co., Inc.; Meso Scale Diagnostics, LLC.; NeuroRx Research; Neurotrack Technologies; Novartis Pharmaceuticals Corporation; Pfizer Inc.; Piramal Imaging; Servier; Takeda Pharmaceutical Company; and Transition Therapeutics. The Canadian Institutes of Health Research is providing funds to support ADNI clinical sites in Canada. Private sector contributions are facilitated by the Foundation for the National Institutes of Health (www.fnih.org). The grantee organization is the Northern California Institute for Research and Education, and the study is coordinated by the Alzheimer’s Disease Cooperative Study at the University of California, San Diego. ADNI data are disseminated by the Laboratory for Neuro Imaging at the University of Southern California.

### Epigenetic Clock and Cell Type Deconvolution

Published epigenetic clocks were calculated according to published methods from processed DNA methylation data. To calculate the principal component-based epigenetic clock for the Horvath multi-tissue clock, Hannum clock, DNAmPhenoAge clock, GrimAge clock, and telomere length we used the custom R script available via GitHub (https://github.com/MorganLevineLab/PC-Clocks). Non-principal component-based (non-PC) Horvath, Hannum, and DNAmPhenoAge epigenetic metrics were calculated using the *methyAge* function in the ENMix R package. The pace of aging clock, DunedinPACE, was calculated using the *PACEProjector* function from the DunedinPACE package available via GitHub (https://github.com/danbelsky/DunedinPACE). We used a 12 cell immune deconvolution method to estimate cell type proportions(*99*).

## Data availability

Anonymized data will be shared by request from a qualified academic investigator for the sole purpose of replicating procedures and results presented in the article.

## Code Availability

All models were built using publicly available packages and functions in the R programming language.

## Acknowledgments Funding

National Institutes of Health grant UM1AI164559 (LCN, DN, MC) National Institutes of Health grant R56 AG078970 (DN), National Institutes of Health grant R01DA052027 (LCN, DN)

## Author contributions

Conceptualization: LCN, RS, MJC Methodology: MB, APP, MJC, NC, VD

Investigation: LCN, MB, VD, APP, RS, MJC, NC, VD Supervision: LCN, MJC

Writing—original draft: MJC

Writing—review & editing: LCN, MJC, ND, DN

