## Supplementary figures and images for "Retroelement-Age Clocks: Epigenetic Age Captured by Human Endogenous Retrovirus and LINE-1 DNA methylation states"

### Figures

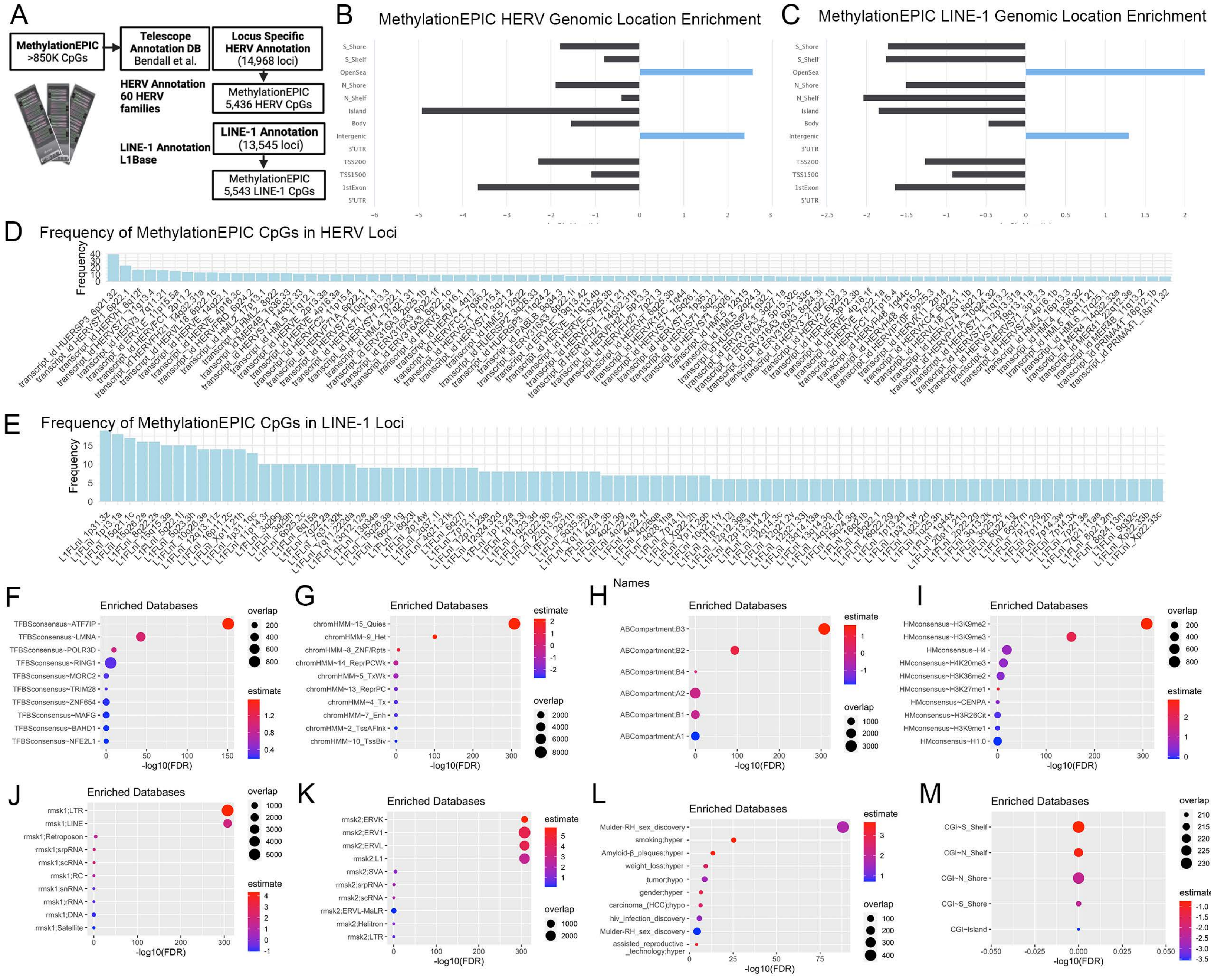

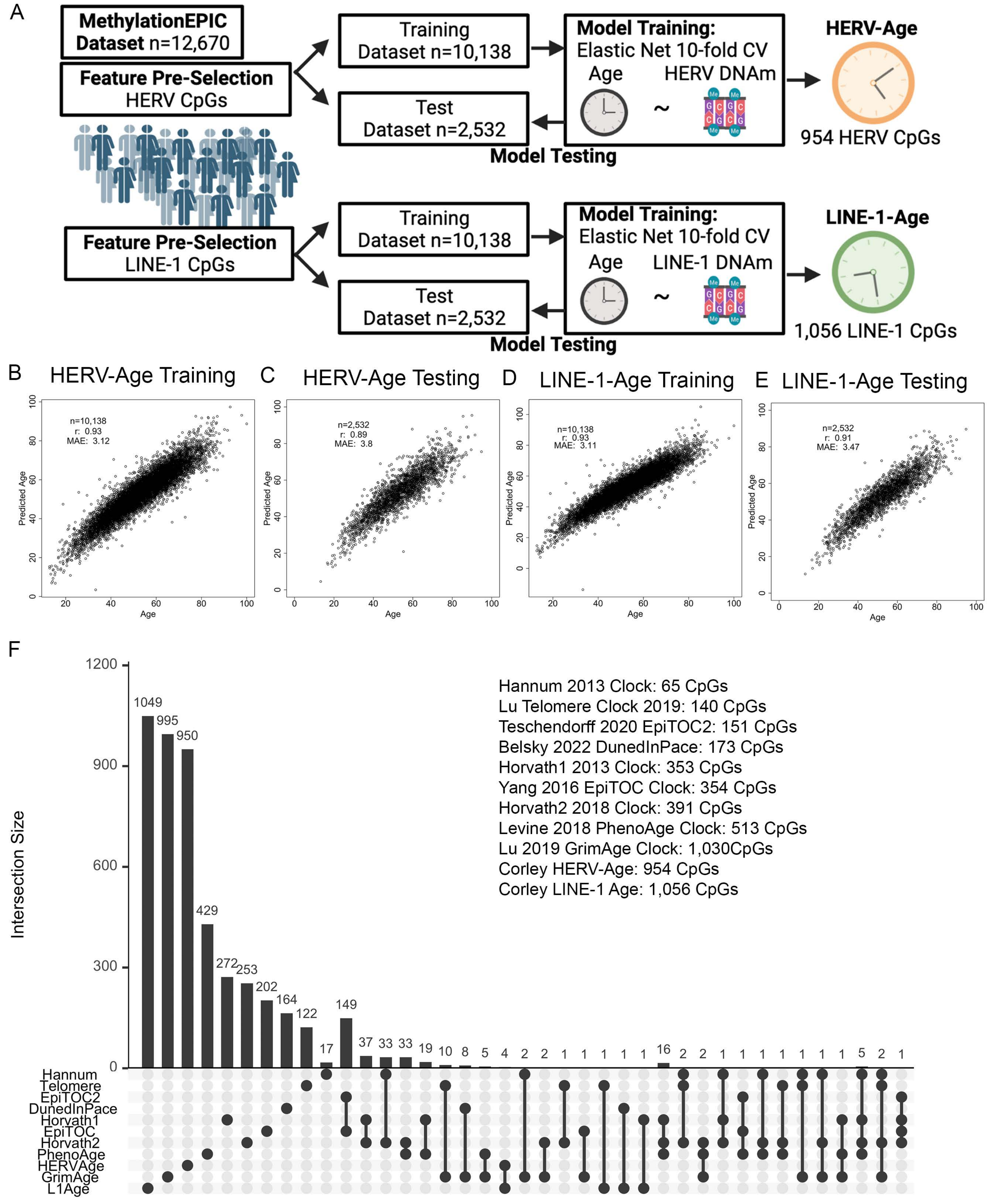

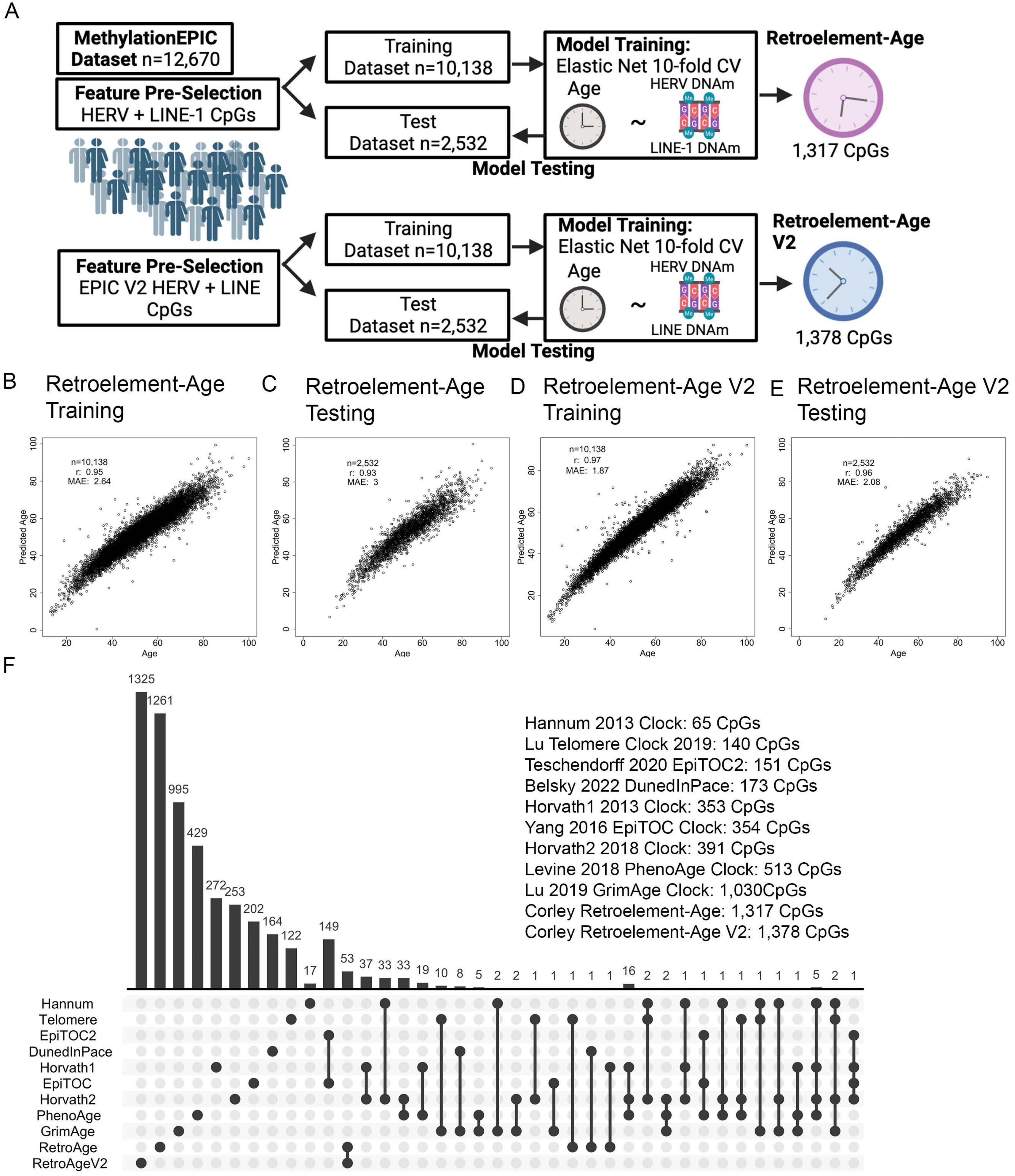

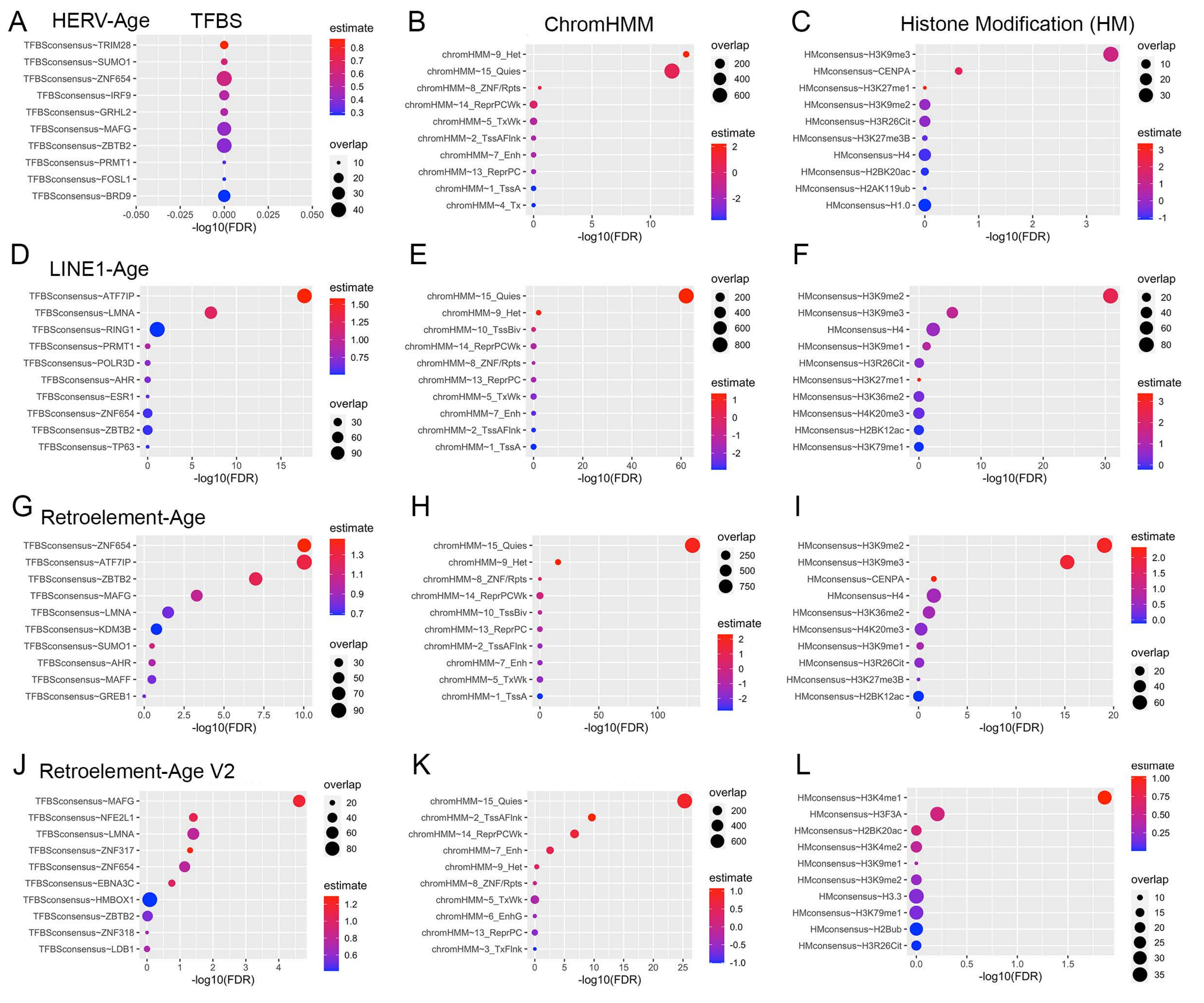

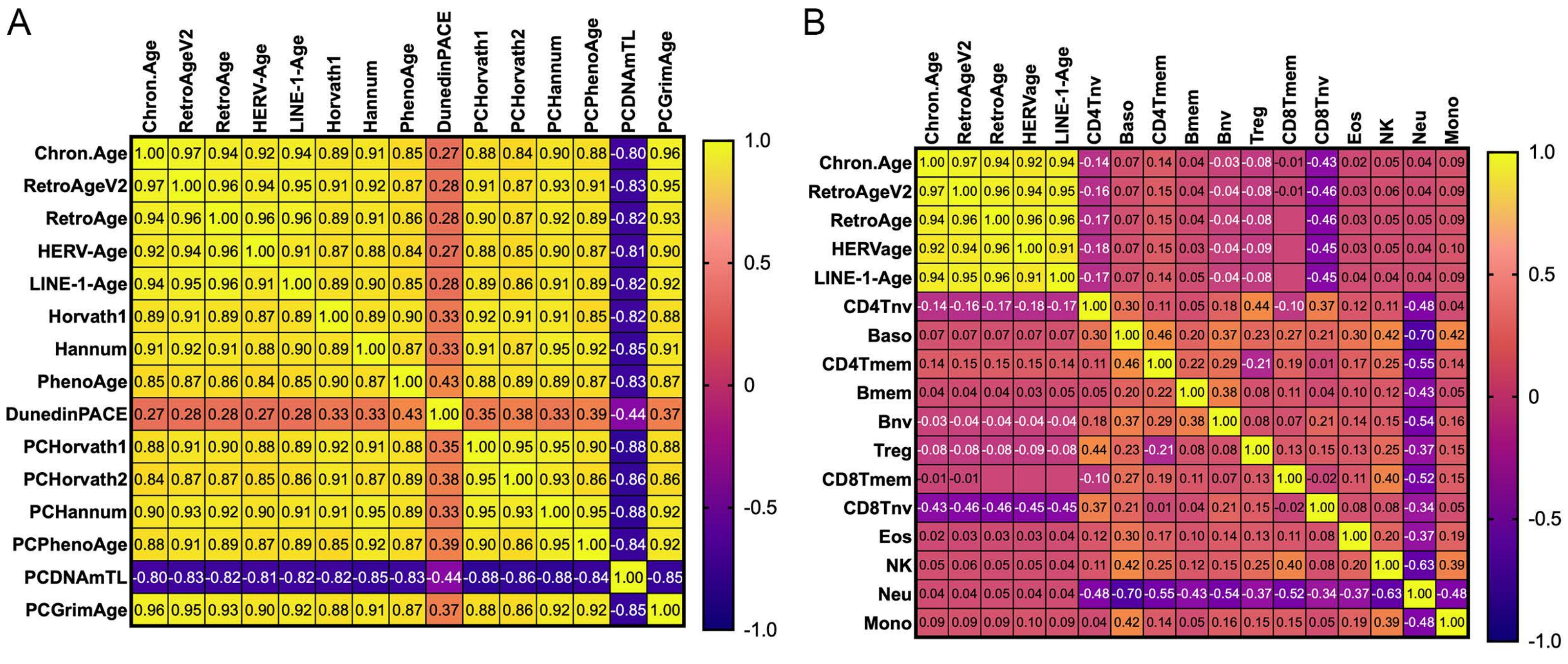

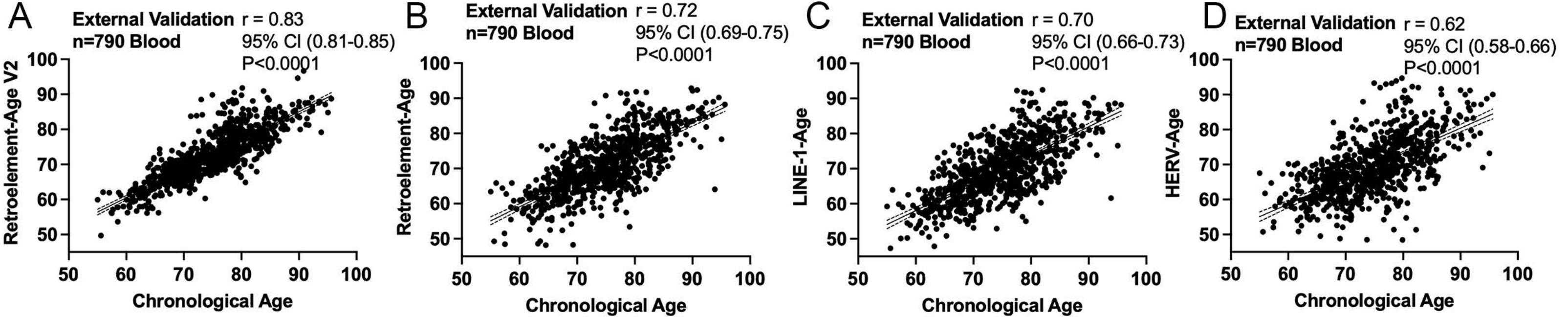

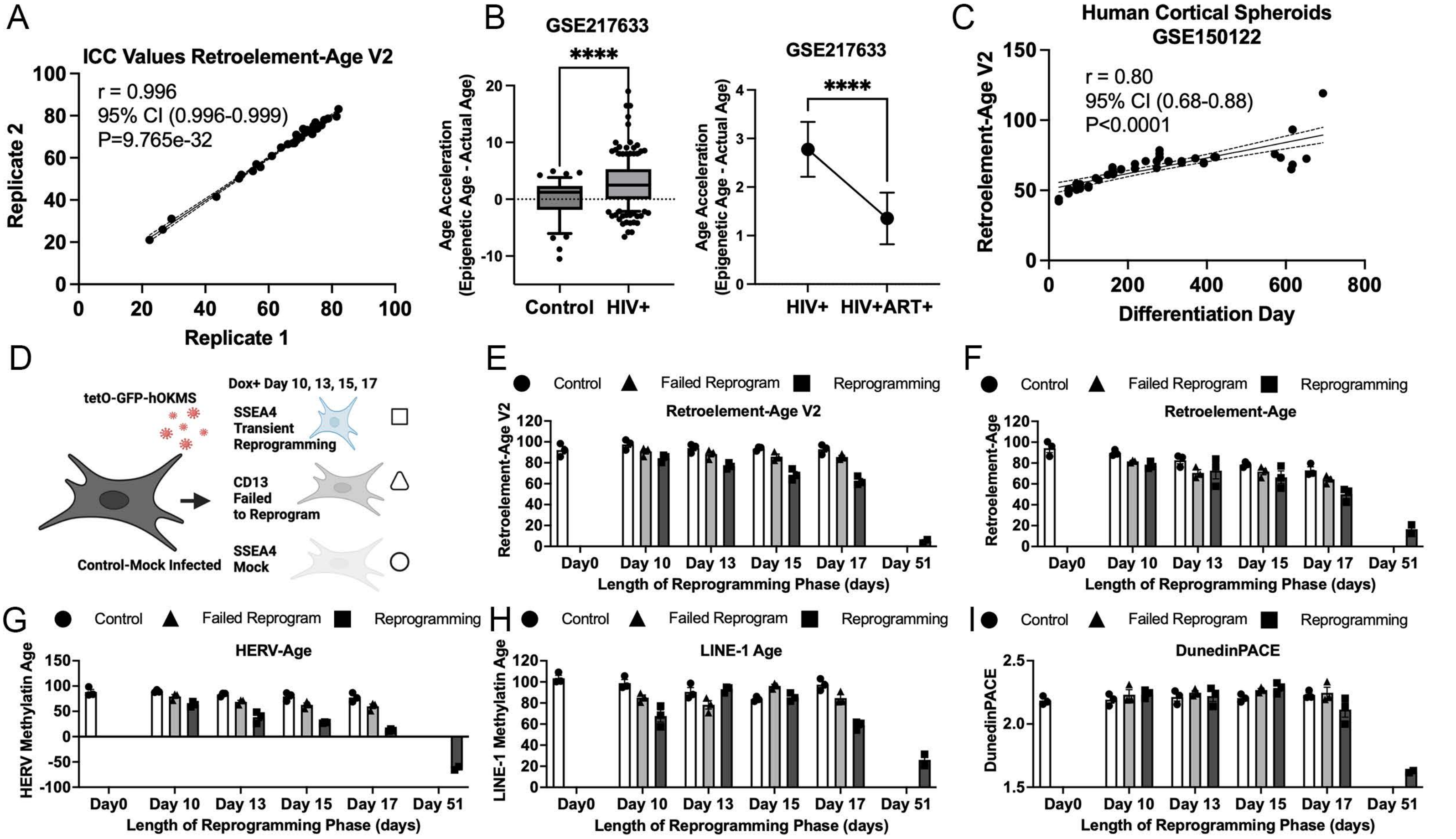

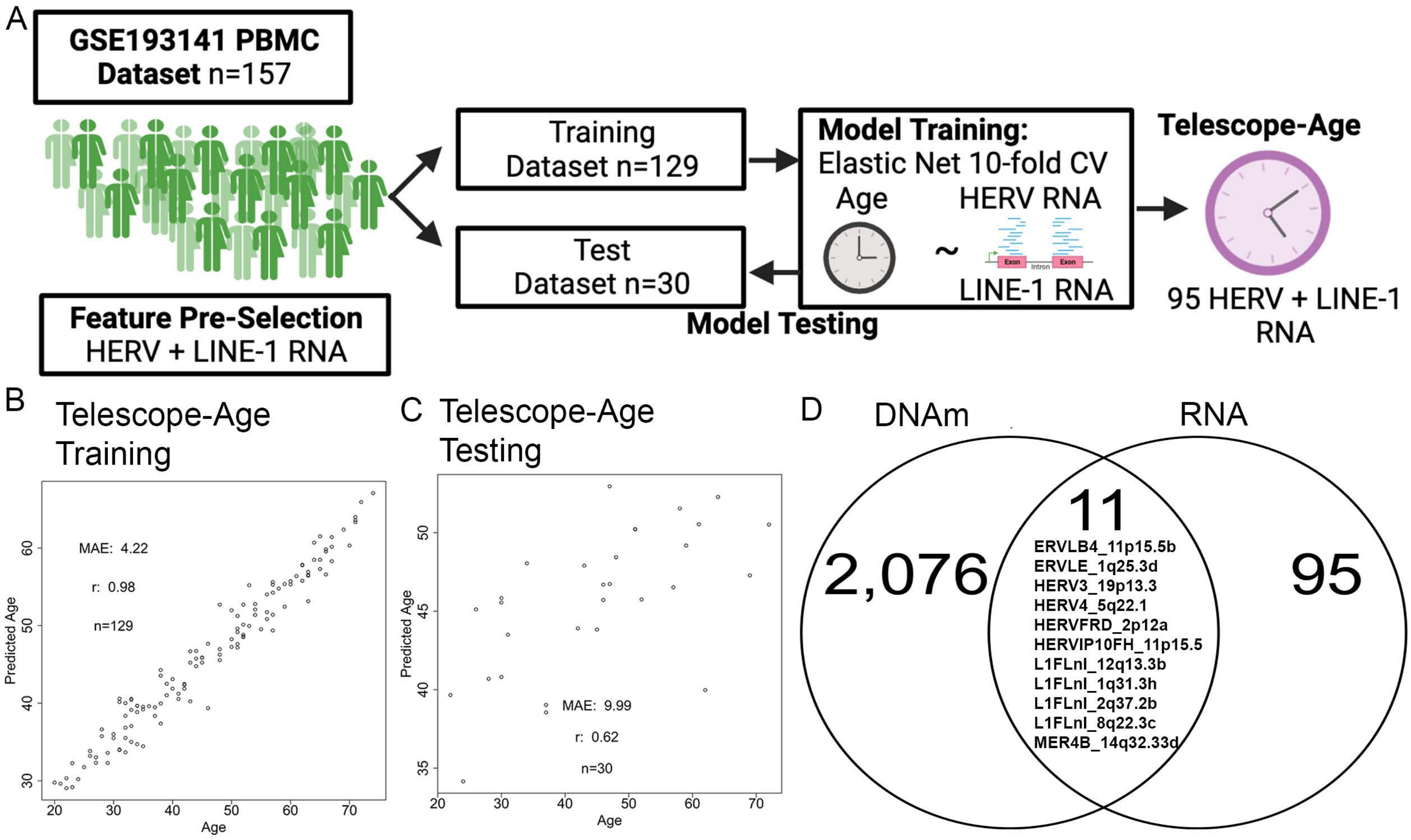

A Multi-Tissue RetroAge Training

B Multi-Tissue RetroAge Testing

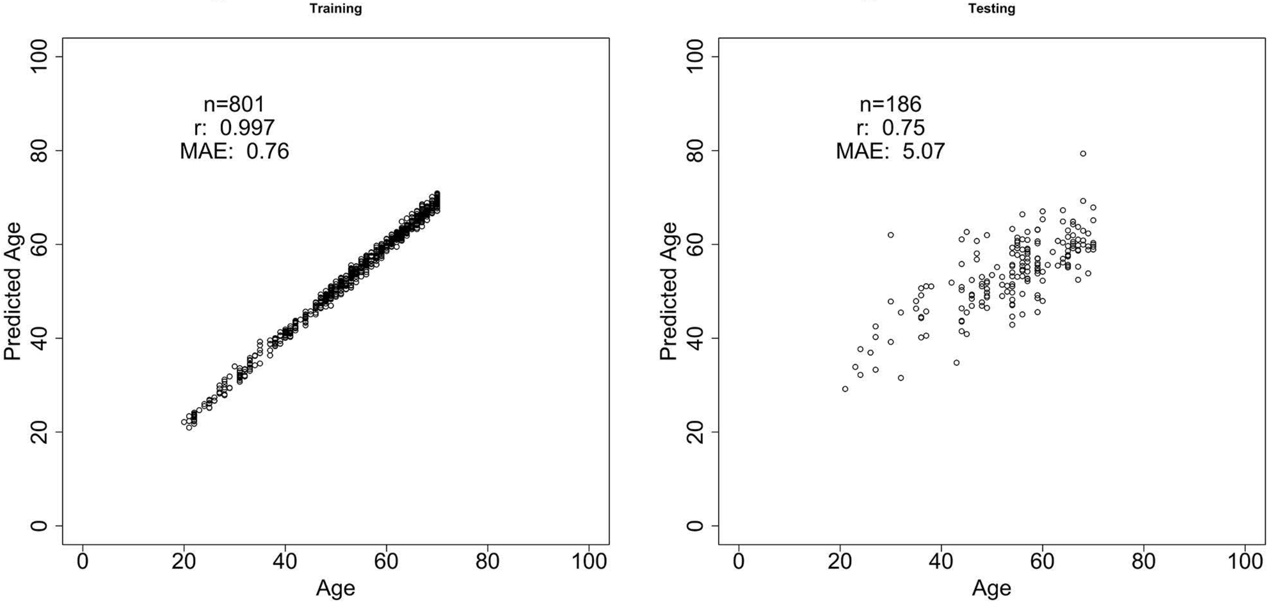

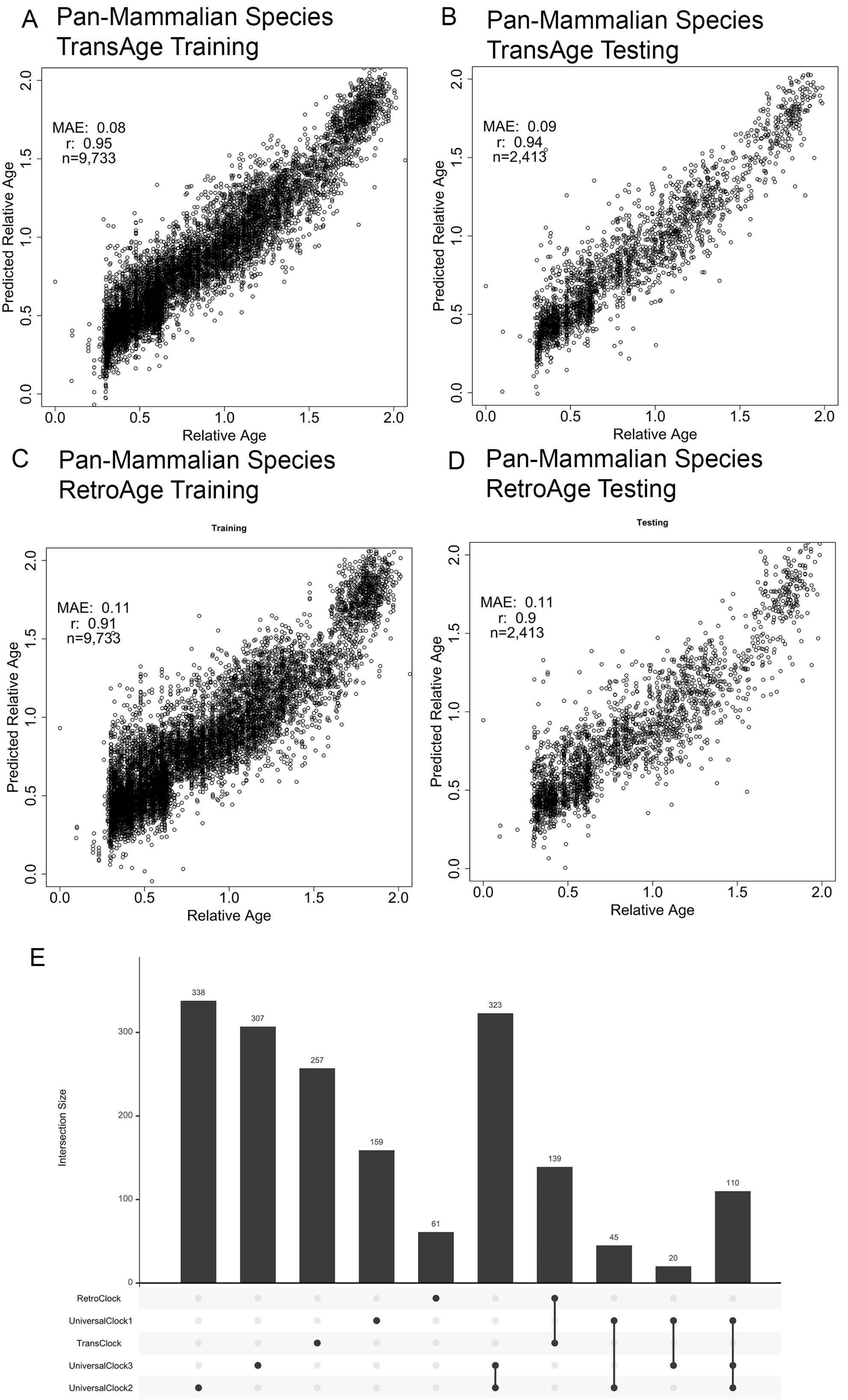
